# A stimulus–state geometry in somatosensory cortex reorganizes during inflammatory pain

**DOI:** 10.64898/2026.08.11.744145

**Authors:** Ara Schorscher-Petcu, Isobel Parkes, Liam E. Browne

## Abstract

Adaptive behavior requires external input to be interpreted together with internal state and ongoing behavior. During pain, noxious somatosensory input evokes movement and arousal, and injury reshapes this relationship, yet how cortical activity organizes stimulus content with behavioral state remains unclear. In awake mice, we delivered hindpaw stimuli while tracking movement, arousal and facial expression, and studied primary somatosensory cortex (S1) using widefield and two-photon calcium imaging, single-action-potential activation of nociceptors, and S1 silencing. Here, we show that S1 neurons were broadly recruited by stimulus and state, whereas latent population dimensions carried mechanical stimulus content. Inflammatory injury caused a reorganization of S1 geometry, binding state and protective responses tighter together. Noxious heat drove S1 as strongly, but engaged mostly the state axis. S1 silencing reduced mechanical hypersensitivity, arousal, and facial expressions. S1 thus embeds mechanical input within a stimulus–state geometry, which is reorganized during inflammatory injury to support adaptation of a coordinated protective response.

## Introduction

Adaptive behavior requires the brain to interpret external input in the context of internal state and ongoing behavior, while preserving information about both. Noxious somatosensory input brings these signals together. It alters movement, arousal, facial expression, and other ongoing behavior, and injury changes how subsequent input is coupled to behavior. How cortical activity preserves stimulus information within ongoing changes in movement and behavioral state remains unknown. Resolving this organization is necessary to interpret cortical activity during pain, where changes in activity could reflect stimulus content, behavioral state, or the way they are embedded together.

Primary somatosensory cortex (S1) is a natural place to study how these signals are organized. S1 is classically viewed as a somatotopic map of peripheral input, but activity in awake S1 is also shaped by movement, arousal, and body configuration (Constantinople and Bruno, 2011; Ferezou et al., 2007; Gantar et al., 2025; Mathis et al., 2017; Musall et al., 2019; Palacio-Manzano et al., 2026; Peron et al., 2015). During nociceptive processing, however, it remains unclear how stimulus, movement, and state signals coexist within the same population. They could be carried by neurons selective for particular inputs, mixed within individual neurons, or distributed across population dimensions from which stimulus information remains recoverable. Resolving this organization requires neural activity and multiple behavioral measures to be recorded together on individual trials.

Knowing how these signals are organized would inform understanding of pain after injury. Inflammation and nerve injury produce marked changes in S1, and manipulating S1 can alter mechanical hypersensitivity and protective behavior (Cichon et al., 2017; Eto et al., 2011; Kim and Nabekura, 2011; Liu et al., 2018; Okada et al., 2021). These findings establish a contribution of S1 without explaining how that contribution changes after injury. Increased responses could reflect greater gain within an otherwise stable population organization, whereas altered population structure could change how somatosensory input is related to movement and state. Distinguishing these possibilities is necessary to understand how cortical processing adapts after injury and why S1 influences some components of the sensitized response but not others.

Here, we delivered somatosensory stimuli in mice while measuring behavior and monitoring or manipulating hindlimb S1 activity. Mechanical, thermal, and transdermal optogenetic stimuli were delivered to the hindpaw, with the latter evoking a single action potential volley in genetically defined nociceptive afferents (Browne et al., 2017). We quantified how widespread behavioral responses covaried across stimuli and trials, sampling local limb movement, broader state-related changes in whisker and pupil dynamics, and facial expressions (Cazettes et al., 2025; Dolensek et al., 2020; Poulet and Petersen, 2008; Reimer et al., 2016). We then asked how this behavioral covariance was reflected in bulk macroscopic and single-neuron S1 activity, and whether stimulus information was carried by selective neurons or latent population dimensions. Finally, we tested how CFA-induced inflammation changed this organization and whether S1 activity was required for sensitized paw and state-related responses. Mechanical stimulus information remained embedded within population activity dominated by movement and state, and inflammatory injury reshaped this organization such that S1 activity became necessary for the coordinated expression of mechanical hypersensitivity, hyperarousal, and facial state.

## Results

### Multi-modal somatosensory stimulation with simultaneous behavioral and neural recordings

To determine how S1 activity relates to somatosensory stimuli, movement, and behavioral state, we developed a setup that combined controlled hindpaw stimulation with simultaneous cortical imaging and high-speed behavioral recording. Head-fixed mice could move their limbs and trunk on a freely rotating grid while the plantar hindpaw remained accessible from below (Fig. 1A,B). This arrangement allowed stimuli, neural activity, and multiple state-related behavioral measures to be aligned on individual trials.

**Figure 1.**
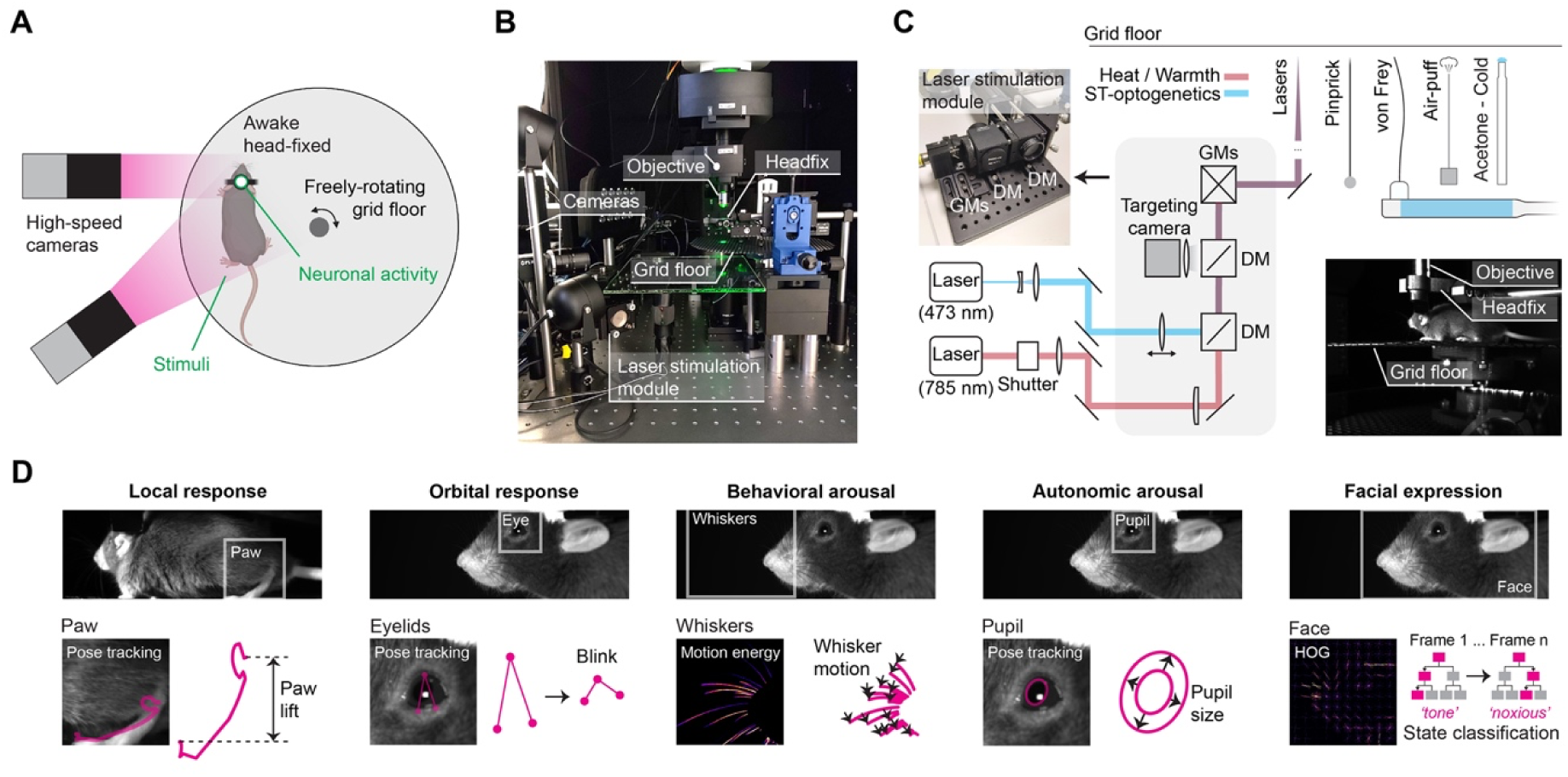
Multi-modal somatosensory stimulation with simultaneous behavioral and neural recordings. (**A**) Schematic of the head-fixed recording configuration. A mouse is head-fixed on a freely rotating grid floor, allowing behavior. Two high-speed machine-vision cameras image the face and body. Somatosensory stimuli are delivered to the plantar surface of the hindpaw through the grid and neural activity is acquired from S1 through the optical access above the headplate (*green arrows*). (**B**) Image of the recording setup showing the two high-speed cameras, the imaging objective (widefield or two-photon), the headfix clamp, the rotating grid floor, and the laser stimulation module below the grid. (**C**) Optical layout of the laser stimulation module (*top left*). A 473 nm laser (*blue path*) provides scanned transdermal (ST) optogenetic stimulation of TRPV1::ChR2 glabrous skin nociceptors; a 785 nm laser (*red path*) with electronic shutter delivers thermal ‘heat’ (10.1 W) and thermal ‘warmth’ (4.1 W) stimuli. Both beams pass through two dichroic mirrors (DM) and are steered by two silver galvanometer mirrors (GMs) to a common targeting path, which directs reflected light back to a targeting camera for hindpaw targeting. ST-optogenetic and thermal laser spot sizes at the paw are 1.25 mm and 4.5 mm, respectively. Other stimuli (blunt insect pin ‘pinprick’; 0.4 g filament ‘von Frey’; 50 psi ‘air-puff’; acetone ‘cold’) and a side-view frame of the awake head-fixed mouse on the grid floor (*bottom right*) are shown. (**D**) Behavioral readouts are extracted from the high-speed video. Top row: representative video frames from the body and face cameras, with the region analyzed in each case highlighted. Bottom row: the specific measurement obtained from each region. ‘Local response’ used paw-lift kinematics (vertical displacement ‘paw lift’ and Euclidean paw displacement ‘paw distance’ with DeepLabCut keypoints). ‘Orbital response’ used eyelid aperture from three keypoints, decomposed into blink, orbital tightening, and orbital dilation. ‘Behavioral arousal’ used whisker motion energy calculated as frame-to-frame absolute pixel difference within the whisker-pad ROI. ‘Autonomic arousal’ used pupil size calculated as an ellipse fit to keypoints. ‘Facial expression state’ used per-frame histogram of oriented gradients (HOG) descriptors and passed to a random forest classifier for per-frame probabilities of pre-defined stimulus-specific categories (tone face, noxious face; see *Methods*).

Through the grid, we delivered a panel of mechanical, thermal, and optogenetic stimuli to the plantar hindpaw (Fig. 1C). Pin-prick, von Frey, and cold stimuli were applied manually, with delivery cued to the experimenter and synchronized to acquisition, whereas warmth, noxious heat, and air-puff were delivered automatically. The laser stimulation module also enabled scanned transdermal optogenetics (ST-opto), allowing selective activation of genetically defined nociceptive afferents with high spatial and temporal precision (Browne et al., 2017; Parkes et al., 2026; Schorscher-Petcu et al., 2021).

Somatosensory stimulation can recruit local movement alongside broader state-related changes. We therefore recorded several behavioral features simultaneously at high speed (Fig. 1A,D). Paw movement, eyelid responses, and pupil size were tracked frame by frame using DeepLabCut (Mathis et al., 2018), whisker movement was quantified from motion energy, and facial features were measured using histograms of oriented gradients (Dolensek et al., 2020). These measures spanned movement, arousal, and facial expression. Recording them together allowed us to examine whether somatosensory stimuli recruited a common behavioral response or different combinations of behavioral features.

We then related these behavioral measures to cortical activity by recording calcium signals in contralateral hindlimb S1 using widefield imaging through a clear-skull preparation (Fig. 3), or two-photon imaging through a cranial window at single-neuron resolution (Fig. 4,5). We also used clear-skull optogenetic inhibition to test whether S1 was required for the coordinated sensory and state response during injury (Fig. 6). Together, these complementary approaches allowed us to determine how stimulus- and state-related signals were organized in S1 and whether S1 activity was required for the accompanying behavioral state responses.

### Somatosensory stimuli recruit variable combinations of behavioral features

Using this platform, we first characterized the multicomponent behavioral states accompanying somatosensory stimulation. Mean responses defined the characteristic profile recruited by each stimulus, whereas trial-by-trial covariance defined how behavioral features were co-expressed within each stimulus condition. Together, these analyses distinguished stimulus-specific from shared behavioral states, and stereotyped responses from variable combinations of features.

Mean responses revealed that each stimulus produced a different mixture of paw movement, whisking, pupil dynamics, and facial expression (Fig. 2A). Noxious heat and pin-prick generally produced larger and faster responses than warmth and von Frey, respectively, while the non- noxious but salient air-puff also recruited several behavioral features (Fig. S1,S2). Noxious stimuli produced a characteristic facial expression, which alone predicted stimulus category above chance (Fig. S3).

**Figure 2.**
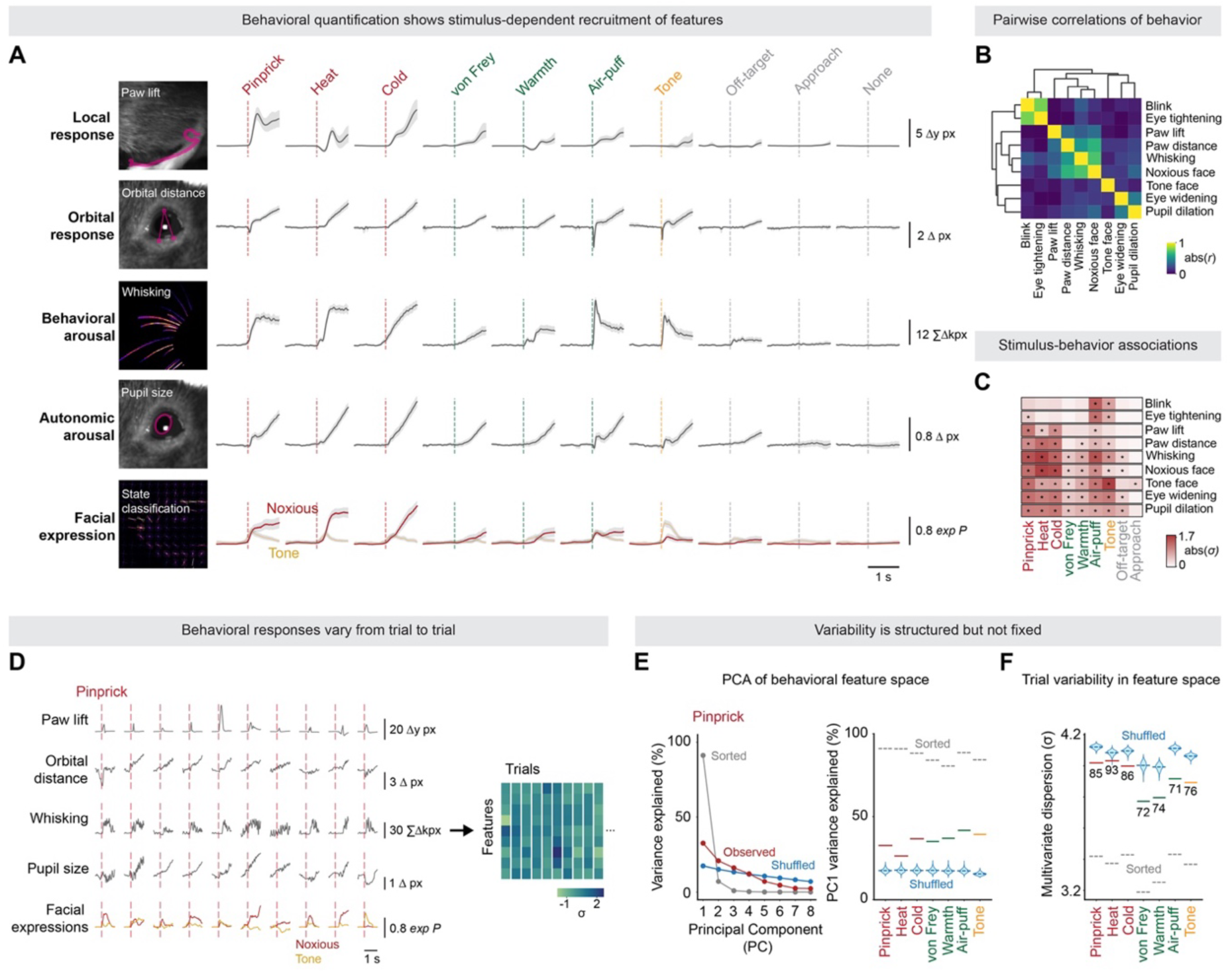
Somatosensory stimuli recruit variable combinations of behavioral features. **(A)** Mean trial- averaged traces for five behavioral metrics (rows) across ten stimulus types (columns). Traces were first averaged across trials within each mouse, then averaged across mice; shading indicates 95% bootstrap CI across mice (n = 6–8 mice; ∼80–120 trials per stimulus in total, with ∼10–18 trials per stimulus for each mouse). *Dashed vertical line* is the stimulus onset with stimulus class indicated by color. (**B**) Pairwise trial-by- trial correlation matrix of the nine behavioral scalars. For each mouse, correlations were determined across trials after pooling all stimuli, then averaged across mice (*n* = 8). Values are absolute Pearson correlations (|*r*|). Dendrograms show Ward-linkage hierarchical clustering on correlation distance. (**C**) Linear mixed-effects analysis of stimulus-behavior associations across the nine behavioral scalars. For each feature, trial-level scalar amplitudes were modeled as *feature ∼ Stimuli + (1 | Mouse).* Heatmap values show the mean per-trial standardized response amplitude for each stimulus-feature pair, after *z*-scoring each feature across stimuli within mouse and then averaging across mice. Asterisks mark cells with BH-FDR-corrected *p* < 0.05 within- feature. **(D)** Individual-trial traces for ten representative pinprick trials (one column per trial), showing paw lift, orbital distance, whisking, pupil size, and the classifier-derived noxious-face (*red*) and tone-face (*orange*) facial expression probability traces. Dashed vertical line, stimulus onset. The right panel shows how for each trial, a scalar is extracted from each behavioral row, producing a per-trial feature vector (*right*, *z*-scored features × trials matrix) **(E)** *Left*: variance explained by each of the first 8 principal components (of a trial-wise PCA in 9- dimensional feature space, fit per stimulus) for pinprick trials (*red*). *Grey* shows a sorted null in which each feature was independently rank-ordered across trials, and *blue* shows a shuffled null in which each feature was independently permuted across trials (1,000 iterations). *Right*: PC1 variance explained per stimulus (*colored bars*) plotted against the sorted-null upper bound (*grey*) and the shuffled-null distribution (*blue violin*, 1,000 iterations). PC1 captures less than the sorted null and more than the shuffled null in every stimulus, indicating that stimuli evoke combinations of behavioral features rather than a single unified response. **(F)** Multivariate dispersion per stimulus, measured as the mean pairwise Euclidean distance (*σ*-units) across *z*- scored feature vectors. Observed dispersion per stimulus is shown (*colored bars*) alongside sorted maximum- structure null (*grey*) and shuffled null (*blue violins*, 1,000 iterations). Numerical annotations give the observed percentile within the shuffled-null distribution. Across stimuli, observed dispersion sits between the sorted and shuffled controls.

Do these features vary together across trials? After pooling stimuli within each mouse, pairwise correlations were calculated between features on individual trials (Fig. 2B,S4; Fig. S5 for analyses within each stimulus). Correlations between features were modest. Strong correlations included those between blink and eye tightening, and between whisking and the noxious face, so some of this covariance may reflect shared motor or anatomical coupling. Some features were coupled therefore, but not all of them together.

Next, linear mixed-effects models identified the features reliably recruited by each stimulus (Fig. 2C). Noxious and other salient stimuli engaged several features, with the greatest overlap between the profiles evoked by noxious stimuli. Nevertheless, different stimuli produced characteristic mean behavioral profiles. These averages concealed substantial variation between individual trials.

On single trials, the same stimulus produced different combinations of behavioral features from one presentation to the next. This variation is illustrated for five features across ten pin-prick trials (Fig. 2D; corresponding von Frey trials shown in Fig. S6). For quantitative analysis, each of nine behavioral features was *z*-scored across trials and combined into a feature vector for each trial. The resulting feature-by-trial matrix revealed pronounced trial-to-trial variation in the behavioral response.

Finally, we asked whether the features varied together across trials or varied independently of one another. Principal component analysis was applied to the nine behavioral measures, separately for each stimulus (Fig. 2E). The variance explained by the leading components was compared with two references: trials in which the correspondence between features was randomly shuffled, and trials in which each feature was rank-ordered to maximize shared variance. Across stimuli, the first principal component explained more variance than the shuffled reference but substantially less than the rank-ordered upper bound. Multivariate dispersion analysis gave the same general result, with the observed values falling between the shuffled and rank-ordered references across stimuli (Fig. 2F). Behavioral features therefore shared structured covariance but did not form a fixed response ensemble.

Noxious and innocuous stimuli thus recruited variable combinations of behavioral features rather than fixed responses. Because the same stimulus was accompanied by different behavioral states on different trials, and these varied rather than moving together as a fixed response, stimulus-averaged neural responses could mix signals related to stimulus identity with those related to the state expressed on each trial. Resolving the two required analyzing S1 activity trial by trial, which we turn to next.

### S1 activity jointly carries mechanosensory, movement, and state signals

Widefield calcium imaging gave access to a large field of view (4.3 × 3.2 mm) through a clear- skull preparation in mice expressing GCaMP6s in excitatory forebrain neurons (Camk2a- tTA::tetO-GCaMP6s). This allowed stimulus-evoked activity to be recorded across contralateral somatosensory cortex in awake mice and under light anesthesia (Fig. 3A). Frames of peak activity showed the spatial distribution of activity across the field of view (Fig. 3B), while mean fluorescence traces showed the time course of the response (Fig. 3C, S7). Anesthesia removed behavioral state signals such as movement and arousal and was used to localize the peak of stimulus-evoked activity to the expected hindlimb region (Fig. 3A,E), defining the region of interest for subsequent trial-level analysis.

**Figure 3.**
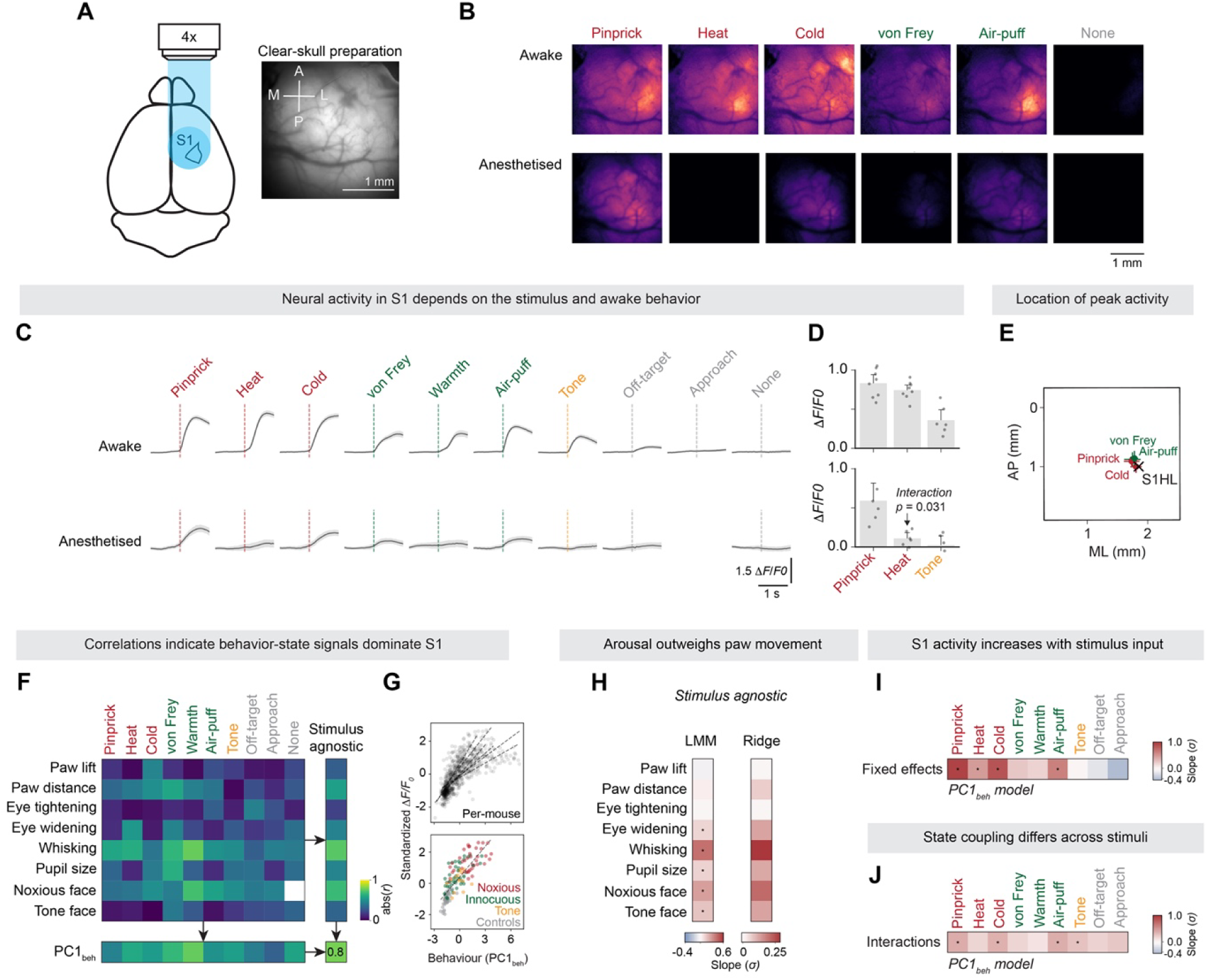
S1 activity jointly carries mechanosensory, movement, and state signals. **(A)** Schematic of the clear-skull widefield preparation in GCaMP6s-expressing mice. *Right*, an example baseline field-of-view. **(B)** Representative stimulus-evoked pixel-wise Δ*F*/*F*₀ peak response maps from one mouse while awake (*top*) or anesthetized (*bottom*). For each stimulus, trial Δ*F*/*F*₀ stacks were averaged frame-by-frame, and the displayed image is the frame with the largest mean fluorescence across pixels. **(C)** Mean Δ*F*/*F*₀ time-courses from the functionally localized S1HL ROI defined in (E), shown for each stimulus in awake (*top*) and anesthetized (*bottom*) sessions. Traces were averaged within mouse and then across mice with 95% bootstrap CI (*n* = 6 mice). **(D)** Scalar Δ*F*/*F*₀ responses, measured as the mean over 0–500 ms post-stimulus, for pinprick and heat, and non-somatosensory tone. Bars show mean across mice, dots show individual mice with 95% bootstrap CI. The awake-to-anesthesia reduction was significantly greater for heat than pinprick (paired exact sign-flip permutation test interaction, *p* = 0.0312). **(E)** Peak locations of stimulus-evoked widefield activity for four localizing stimuli in anesthetized mice: pinprick, cold, von Frey, and air-puff. Crosses show the mean ± SEM peak coordinate across mice (*n* = 6). The black cross marks the stereotaxic coordinate of S1HL (−0.70 mm AP, 1.85 mm ML relative to Bregma). Axes show offset from Bregma. **(F)** Pairwise absolute Pearson correlations between S1 Δ*F*/*F*₀ and each behavioral scalar, determined trial-by-trial within each stimulus and awake mouse (*n* = 6). S1 activity was measured as the mean Δ*F*/*F*₀ over 0–500 ms in the functionally localized S1HL ROI. The rightmost column shows stimulus-agnostic correlations calculated after pooling trials across stimuli within mouse. The bottom row shows correlations between S1 activity and PC1 of the eight-feature behavioral matrix (PC1_beh_). **(G)** Scatter of standardized S1HL Δ*F*/*F*₀ as a function of PC1_beh_ score. *Top*, all trials pooled across mice with dashed lines showing separate linear fits for each mouse (*n* = 6). Bottom, trials from one representative mouse colored by stimulus identity, with the *dashed line* showing the linear fit for that mouse. (**H**) Stimulus-agnostic models relating S1HL Δ*F*/*F*₀ to behavioral features across all trials. *Left*, LMM fixed-effect slopes from *activity ∼ behavioral predictors +* (*1 | Mouse*), where behavioral predictors consisted of the eight individual behavioral scalars*. Right*, ridge-regression coefficients using the same predictors, fit separately for each mouse with 10-fold cross-validation. (**I**) Per-stimulus fixed effects from the model *activity* ∼ *PC1_beh_ × Stimuli* + (*1* │ *Mouse*). Values show the shift in S1HL Δ*F*/*F*₀ for each stimulus relative to the no-stim reference, at mean behavioral state. (**J**) Interaction terms from the same model, showing how far each stimulus-specific *PC1_beh_* slope departs from the stimulus-agnostic slope. Non-zero values indicate that the relationship between behavioral state and S1 activity depended on the stimulus. Asterisks mark BH-FDR- corrected p < 0.05.

Mechanical and cold responses largely persisted under anesthesia, whereas heat and tone responses were abolished (Fig. 3C,D, S8). The differential effect of anesthesia suggested that heat- and tone-evoked S1 activity depended strongly on the awake behavioral state. We therefore asked whether S1 activity in awake mice covaried with behavior on individual trials. For each stimulus and behavioral feature, Pearson correlations were calculated between the S1 response magnitude and the corresponding behavioral measure on the same trial (Fig. 3F).

Several features correlated strongly with neural activity, most notably whisking, pupil dynamics, and facial expression. Correlations remained high when computed across stimuli, and when the first principal component of behavior (PC1_beh_) was used in place of individual features (Fig. 3G).

These correlations indicated that a substantial component of macroscopic S1 activity covaried with trial-to-trial behavioral state. Linear mixed-effects models and ridge regression were then used to quantify the contributions of the behavioral features to S1 activity (Fig. 3H; Fig. S9 for analyses within each stimulus). Whisking, pupil dynamics, and facial expression were stronger predictors of neural activity than movement of the stimulated paw, arguing against a simple account based on local movement or reafference.

If behavioral state dominated S1 activity, could a contribution of stimulus identity still be resolved? Models containing PC1_beh_ alongside stimulus terms resolved significant stimulus effects for several stimuli (Fig. 3I), showing that a single behavioral composite did not fully account for S1 activity. Interactions in the same models further showed that the relationship between behavioral state and S1 activity differed across stimuli (Fig. 3J).

Stimulus and behavioral state were therefore mixed in macroscopic S1 activity: mechanical and cold responses persisted under anesthesia while heat did not, and trial-level analyses showed strong covariation with whisking, pupil dynamics, and facial expression in awake mice. Whether these components are resolved at the level of single neurons, or are instead embedded in a population code, required recordings of individual neurons across S1.

### Single S1 neurons respond broadly and are dominated by behavioral state

Two-photon calcium imaging was used to determine how the stimulus- and state-related signals observed with widefield imaging were distributed across individual neurons. Hundreds of neurons were recorded per session, using three sessions per mouse, in a 512 × 512 µm field of view (Fig. 4A), and spiking events were inferred from the calcium signal by deconvolution (Fig. 4B). ST- opto was included to evoke a single-action-potential volley selectively in genetically defined nociceptors (Browne et al., 2017; Parkes et al., 2026; Schorscher-Petcu et al., 2021).

**Figure 4.**
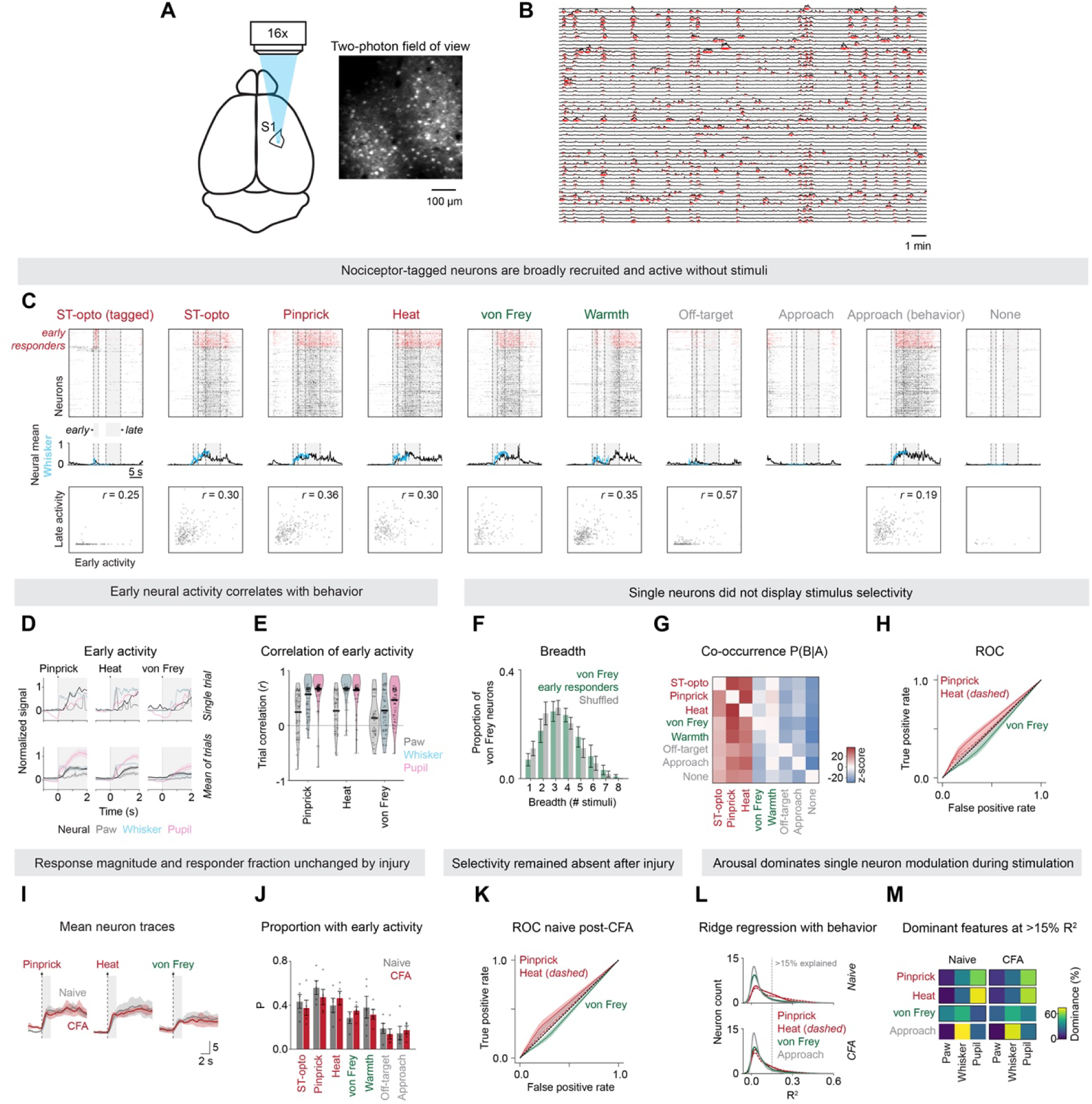
Single S1 neurons respond broadly and are dominated by behavioral state. (**A**) Two-photon imaging schematic and example field-of-view showing GCaMP6s-expressing neurons in hindlimb S1. (**B**) Representative Δ*F*/*F*₀ traces from simultaneously recorded neurons in one session. *Red ticks* illustrate suprathreshold events from the deconvolved activity traces used in analysis, thresholded for display. (**C**) Single-neuron event activity across somatosensory stimuli and control conditions from one representative session. *Top*, event rasters ordered by response to ST-optogenetic activation of cutaneous nociceptors, with early ST-opto-responsive neurons (nociceptor-tagged) marked in red. These neurons were also recruited by heat and by approach, despite no applied somatosensory stimulus during approach. *Middle*, population-mean neural event activity overlaid with whisker motion. *Bottom*, early-late activity correlations. Dashed vertical lines mark analysis windows. (**D**) Normalized single-trial and trial-averaged traces showing mean neural event activity, whisker motion, pupil size, and paw distance. Trial-averaged traces were averaged within mouse and then across mice. (**E**) Per-neuron trial-by-trial correlations between early neural activity and behavioral scalars. Violin plots show distributions across neurons, with black lines indicating means. Neural activity was broadly correlated with behavioral state across stimuli. (**F**) Response breadth of von Frey early-responsive neurons. Breadth is the number of stimuli for which each neuron was classified as an early responder. Colored bars show the observed distribution and grey bars show a matched random-neuron null that preserves all other stimulus responses. (**G**) Co-occurrence of early-responsive populations across stimulus categories. Values show P(B|A), *z*-scored against a shuffle null that preserves each neuron’s response breadth but randomly reassigns stimulus labels. Red indicates greater-than-null co-occurrence and blue indicates lower-than-null co-occurrence. (**H**) ROC curves, using early single-neuron activity as the classifier score. Curves show mechanical versus thermal, pinprick versus others, von Frey versus others, and heat versus others. Curves remain close to the diagonal, indicating weak single-neuron stimulus modality discrimination. (**I**) Mean Δ*F*/*F*₀ traces before and after CFA. (**J**) Bar plot showing proportion of neurons classified as early-responsive to each stimulus before and after CFA. (**K**) ROC curves after CFA, plotted as in (H). Curves still indicate weak single-neuron stimulus modality discrimination. (**L**) Per-neuron ridge-regression R^2^ values for models predicting neural activity from paw movement, whisking, and pupil size. Models were fit separately for conditions, with predictors and neural activity *z*-scored across trials. Dashed line marks R^2^ = 0.15, the threshold used for dominance analysis in (M). (**M**) Dominant behavioral predictor among neurons with ridge R^2^ > 0.15. Dominance was assigned to the feature with the largest absolute standardized ridge coefficient for each neuron. Color indicates the fraction of neurons dominated by each feature. Pupil and whisker responses dominated more often than paw movement. Five mice, 20 sessions (15 baseline, 5 CFA), 9,995 neurons and 409 stimulus trials in total (268–770 simultaneously recorded neurons and 18–22 trials per session). Mouse is the unit of analysis throughout; per-mouse values are averaged across sessions before statistical testing. Bars and shaded regions show mean ± SEM.

We defined neurons with an early ST-opto response as *nociceptor-tagged* neurons. If S1 contained a dedicated nociceptive population, these neurons should respond preferentially to noxious input and show little recruitment otherwise. Instead, ST-opto evoked distinct early and late response phases (Fig. 4C, S10), and the same neurons were recruited by other stimuli and active during spontaneous behavior in the absence of any somatosensory stimulus (Fig. 4C). Early and late magnitudes correlated moderately across trials (r ≈ 0.3, ∼9% shared variance; Fig. 4C, bottom), so even with a genetically and temporally precise input the early phase could not be cleanly separated from state-related activity.

What signal did the early responses carry? Trial-by-trial analysis showed that pupil and whisker dynamics were the strongest correlates of early responses (Fig. 4D,E). Even at single-neuron resolution, and with selective nociceptor input, early S1 activity covaried strongly with whisking and pupil dilation.

We then used three analyses to test stimulus selectivity directly. Response breadth across stimulus types matched a shuffled null (Fig. 4F, S11). Co-occurrence probabilities showed no preferential overlap between responder populations (Fig. 4G, S12). ROC analysis confirmed that single-neuron responses could not reliably distinguish pin-prick from heat or von Frey, nor mechanical from thermal stimuli (Fig. 4H, S12). Although mechanical responses largely persisted under anesthesia whereas heat responses were abolished, this difference was not reflected in a readily identifiable population of neurons selective for mechanical rather than thermal input.

We next asked whether these properties changed after inflammatory injury. Mice received an intraplantar injection of complete Freund’s adjuvant (CFA) into the left hindpaw, and imaging was repeated once inflammation had developed. We observed no change in response magnitude or kinetics (Fig. 4I, S13), the proportions of early and late responses (Fig. 4J, S10), or stimulus selectivity (Fig. 4K, S11). Ridge regression applied to each neuron showed that pupil dynamics was the strongest predictor of single-neuron activity, with whisking also contributing and paw movement generally weaker, and this was unchanged after CFA (Fig. 4L,M, S14,S15).

Single S1 neurons therefore responded broadly, mostly lacked stimulus selectivity, were dominated by behavioral state, and these properties were unchanged by injury. Stimulus identity was not resolved in individual neurons, leaving open whether it is instead carried in the coordinated activity of the population.

### S1 population geometry organizes stimulus and state signals and is reshaped by inflammation

S1 activity was organized along latent population dimensions. For each session, a single PCA was fitted to the deconvolved activity of simultaneously recorded neurons across all stimuli, and trajectories were then examined per stimulus (Fig. 5A). The first three components differed in their trajectories (Fig. 5B) and in their responses across stimuli (Fig. 5C). The leading component appears to capture the dominant response shared across stimuli, while the second and third carried additional stimulus-dependent structure.

**Figure 5.**
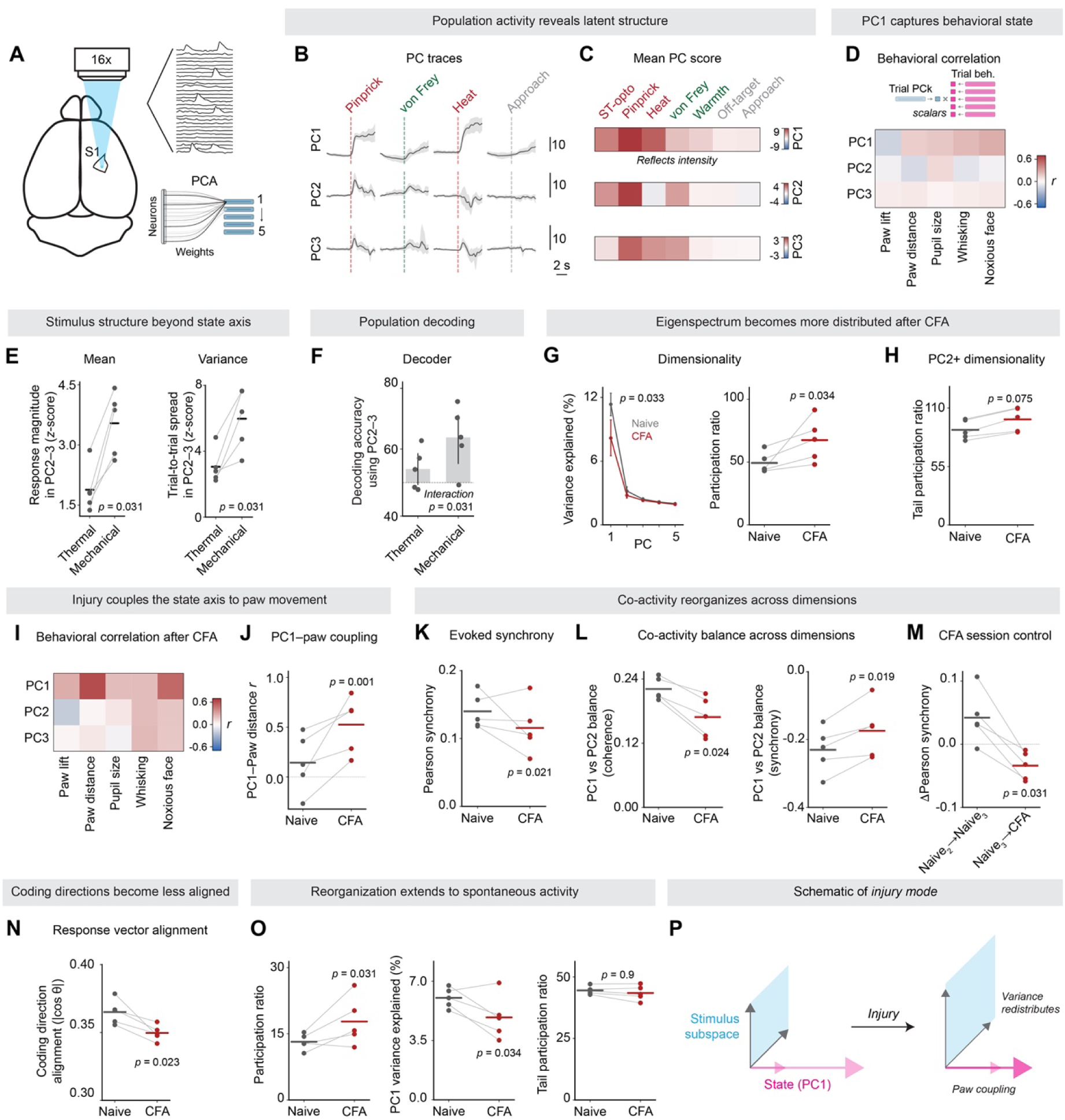
S1 population geometry organizes stimulus and state signals and is reshaped by inflammation. (**A**) Two-photon calcium recordings were deconvolved and projected into PC space to obtain trial-level PC scores and neuron weights. (**B**) Mean PC1–PC3 time courses for representative stimuli. Traces show mean ± SEM across mice. The *dashed line* is stimulus onset. (**C**) Mean baseline-subtracted PC1–PC3 responses for each stimulus, measured over 0 to 1.5 s post-stimulus relative to −1 to 0 s baseline, averaged across mice. (**D**) Stimulus-controlled PC–behavior correlations. To isolate state-related coupling from stimulus-driven differences, PC scores and behavioral scalars were demeaned within each stimulus type before calculating trial-by-trial Pearson correlations, then averaged across mice. (**E**) Response magnitude (left, mean |PC2–3 score| across trials) and trial-to-trial spread (right, standard deviation across trials in PC2–3) for thermal (heat, warmth) and mechanical (pin-prick, von Frey) stimuli. Stimuli were averaged within modality for each mouse. (**F**) Separation of stimuli from control conditions (off-target and approach) using PC2–3 alone, decoded with shrinkage LDA (shrinkage = 0.5) and leave-one-out cross-validation. Mean balanced accuracy across mice with 95% bootstrap CI. Mechanical stimuli were better separated than thermal stimuli in these dimensions. (**G**) Explained variance ratio for the leading PCs and participation ratio of the full signal eigenspectrum before and after CFA. Lines and dots show individual mice, horizontal bars indicate means. (**H**) Tail participation ratio calculated from the signal eigenspectrum after excluding PC1. (**I**) Stimulus-controlled PC–behavior correlations after CFA, calculated as in (D). (**J**) Per-mouse Pearson correlation between PC1 score and paw distance (scalar, stim-controlled). (**K**) Mean pairwise Pearson correlation across all neuron pairs during the evoked window (−2 to +3 s). (**L**) Co-activity balance across dimensions. *Left*, coherence balance (cosine-based d12): the difference in cosine co-activity carried by PC1 versus PC2, capturing both tonic level and temporal fluctuation. *Right*, synchrony balance (Pearson-based d12): the same comparison using Pearson correlation, which isolates temporal co-fluctuation after removing each neuron’s mean. (**M**) The change in Pearson synchrony between adjacent baseline sessions (Naive_2_→Naive_3_) is compared with the transition from the final baseline session to CFA (Naive_3_→CFA). Exact sign-flip permutation (2^5^ = 32 permutations) on the paired session-order difference. (**N**) For each pair of stimuli, a contrast vector was calculated as the normalized difference between stimulus-mean response vectors in neuron space (mean response over 0 to 1.5 s). Values show the mean absolute cosine between all pairs of contrast vectors across five stimuli. Lower values indicate less aligned stimulus contrasts. (**O**) Participation ratio, PC1 explained variance, and tail participation ratio calculated from spontaneous frames outside stimulus-evoked periods. Unless otherwise noted, *p*-values are from exact within-mouse CFA relabeling tests (4^5^ = 1024 permutations, one-tailed). (**P**) Schematic illustrating how variance redistributes across population dimensions and the dominant direction of variation (PC1) couples more tightly to paw movement after injury. Five mice, 20 sessions (15 baseline, 5 CFA), 9,995 neurons and 409 stimulus trials in total (268–770 simultaneously recorded neurons and 18–22 trials per session). Mouse is the unit of analysis throughout; per-mouse values are averaged across sessions before statistical testing.

The leading component tracked behavioral state, correlating with arousal and facial measures across trials of the same stimulus type (Fig. 5D; Fig. S19 without stimulus removal). The second and third components showed no consistent behavioral correlation. Responses in these dimensions were larger and more variable for mechanical than for thermal stimuli (Fig. 5E), and mechanical stimuli were better separated from conditions without hindpaw stimulation than thermal stimuli were, when only these dimensions were used (Fig. 5F). This ordering was not explained by how strongly each stimulus engaged the state axis. Pin-prick and heat both separated strongly from control conditions along the leading component, whereas von Frey did not, even though von Frey was the stimulus best separated in the higher dimensions. Mechanical stimulus content was therefore carried in population dimensions beyond the state axis.

Did inflammatory injury change this population structure? After CFA, the eigenspectrum flattened: the leading components explained less variance and participation ratio (the effective number of dimensions) increased (Fig. 5G,H; S16-18). The leading component also coupled more tightly to paw movement (Fig. 5I,J; S19), so that injury tightened the relationship between the state axis and paw movement.

What underlies this reorganization? We next asked how coordinated activity was distributed across population dimensions. Pearson synchrony captured correlated fluctuations after subtracting each neuron’s within-trial mean, whereas cosine similarity retained the shared offset and was therefore sensitive to population-wide co-activation or coherence. To estimate the influence of each dimension, one component was removed in PC space, the remaining activity was reconstructed in neuron space, and pairwise co-activity was recalculated. The leading component accounted for most coherence, whereas the second and third contributed more strongly to synchrony (Fig. S20,S21). After CFA, total evoked synchrony decreased (Fig. 5K), but the balance across dimensions also shifted: the influence of the leading component on coherence fell, whereas its influence on synchrony increased (Fig. 5L). This redistribution of synchrony and coherence across dimensions was confined to stimulus-evoked activity, was absent during baseline epochs (Fig. S22,S23), and was not explained by session order (Fig. 5M, S23). Inflammatory injury therefore redistributed coordinated activity across population dimensions.

Finally, stimulus contrast became less aligned after injury. For each pair among the five stimuli, a stimulus contrast vector was computed as the normalized difference between stimulus-mean response vectors in neuron space, and their alignment measured as the mean absolute cosine across all pairs (Fig. 5N). Alignment decreased after CFA. This was not explained by the spatial distribution of neuronal loadings (Fig. S18), nor by response magnitude or trial-to-trial noise structure (Fig. S24). CFA-dependent redistribution of variance across the eigenspectrum was also present during spontaneous activity (Fig. 5O), indicating that this aspect of the population reorganization was tonic.

After CFA, single-neuron response magnitude and selectivity remained broadly similar, but the population reorganized (Fig. 5P): variance redistributed across dimensions, the state axis coupled more tightly to paw movement, and stimulus contrasts became less aligned.

### S1 activity is required for mechanical hypersensitivity and hyperarousal after inflammation

Did this reorganization contribute to behavioral state? We silenced S1 during stimulation and asked whether the sensitized response depended on stimulus modality. Right S1HL, contralateral to the injected paw, was inhibited optogenetically in VGAT-ChR2-EYFP mice during interleaved control and inhibition trials time-locked to stimulation. Behavior was measured before and after CFA injection into the contralateral hindpaw (Fig. 6A).

**Figure 6.**
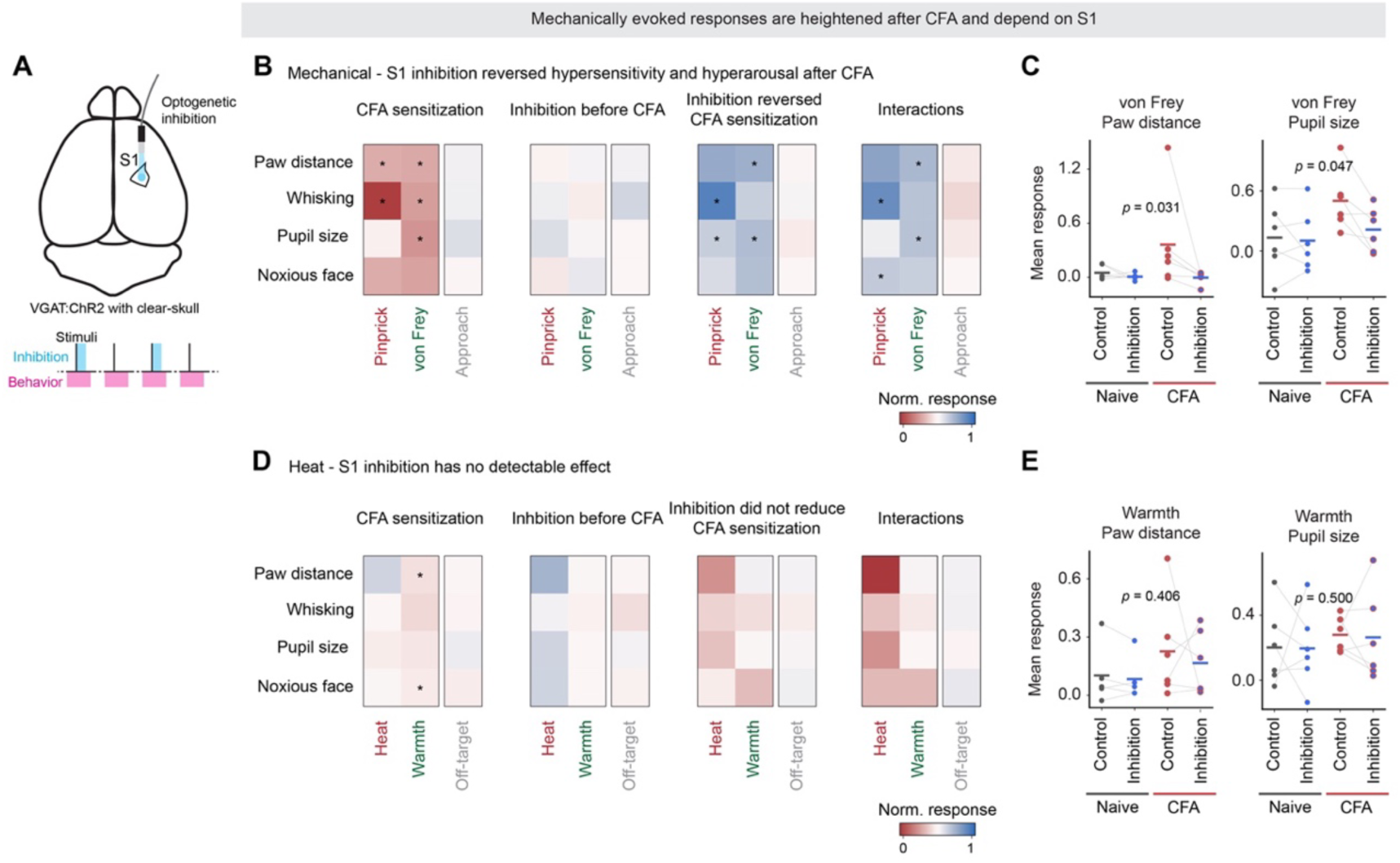
S1 activity is required for mechanical hypersensitivity and hyperarousal after inflammation. (**A**) Optogenetic S1 inhibition strategy. In VGAT-ChR2-EYFP mice a 400 µm-core optical fiber was positioned over hindlimb S1 through a clear-skull preparation. On inhibition trials, 473 nm light was delivered for 1 s at stimulus onset as a 40 Hz sinusoid, with 1.5 mW peak power at the fiber tip and linear attenuation over the final 100 ms. Each stimulus was delivered four times per session: two control and two interleaved inhibition trials. Mice were tested once before and once after CFA. Blue-light masking flashes were delivered randomly throughout to control for off-target visual artifacts. (**B**) Mechanical-stimulus behavioral responses across four contrasts: CFA sensitization, inhibition effect before CFA, inhibition effect after CFA, and interaction between condition and inhibition. Values are normalized by the pooled across-mouse SD for each feature. (**C**) Per-mouse responses for von Frey paw distance and pupil dilation. Asterisks denote exact within-mouse sign-flip permutation tests on the condition × treatment interaction (mouse-level values, one-tailed; * *p* < 0.05). After CFA, S1 inhibition reduced von Frey-evoked paw movement (*p* = 0.031), pin-prick-evoked whisking (*p* = 0.031), and von Frey-evoked pupil responses (*p* = 0.047), and lowered the probability of a von Frey paw response (*p* = 0.015). The two strongest interactions were confirmed by a trial-level linear mixed model (von Frey paw movement, *F*_1,71_ = 8.71, *p* = 0.004; pin-prick whisking, *F*_1,80_ = 4.16, *p* = 0.045); the von Frey pupil interaction was directionally consistent (mixed-model inhibition effect after CFA, *p* = 0.041). S1 inhibition had no significant effect on any measure before CFA, and no interaction reached significance for warmth, heat, or the ipsilateral-hemisphere control (all *p* > 0.15). (**D**) Thermal and control-stimulus responses, plotted as in (B). S1 inhibition did not detectably reduce thermal responses after CFA. (**E**) Per-mouse responses for warmth paw distance and pupil dilation. Six mice, 12 sessions (6 baseline, 6 CFA) and 857 stimulus trials in total.

CFA produced a broad sensitization of stimulus-evoked behavior. Responses generally increased across paw movement, whisking, pupil dilation, and facial expression, during mechanical and warmth stimulation, although the precise profile varied between stimuli, likely reflecting differences in dynamic range (Fig. 6B–E; S25,S26). Heat responses were already large before CFA and changed little. Inflammation therefore extended sensitization beyond the stimulated paw to facial expression and arousal-related pupil and whisker responses. We term this increased stimulus-evoked arousal *hyperarousal*.

S1 inhibition selectively reduced the sensitized mechanical response across paw and state- related measures. Across pin-prick and von Frey stimulation, the effects involved paw movement, pupil dilation, whisking, and facial expression after CFA. Significant CFA × inhibition interactions were found for von Frey-evoked paw displacement and pupil dilation, and for pin- prick-evoked whisking and facial expression (Fig. 6B,C; S25,S26). Ipsilateral S1 inhibition produced no interactions on any measure (all *p* > 0.15), consistent with specificity to contralateral hindlimb S1. S1 inhibition had little effect before CFA. S1 activity was thus required after inflammation for a coordinated mechanical response spanning hypersensitivity, hyperarousal, and facial expression.

Warmth- and heat-evoked responses were not reduced by S1 inhibition after CFA (Fig. 6D,E). This was particularly informative for warmth, which showed sensitization in paw movement and facial expression. Sensitization alone therefore did not confer S1 dependence. If anything, heat- evoked paw responses were larger during inhibition after CFA. This dissociation matched the preceding neural recordings: warmth and heat carried little stimulus structure in the population dimensions beyond the state axis (Fig. 5E,F), and heat-evoked S1 activity was largely abolished under anesthesia (Fig. 3C,D).

The causal dissociation mirrored the organization of stimulus and state signals in S1. Mechanical responses depended on S1, consistent with mechanical stimulus identity being carried in population dimensions alongside state-related activity. By contrast, sensitized warmth- and heat- evoked responses remained intact, consistent with their stimulus identity carrying little such structure. S1 activity was not only correlated with mechanical hypersensitivity and hyperarousal after inflammation, but necessary for their coordinated expression.

## Discussion

S1 responses to noxious stimuli have often been interpreted within a sensory-gain framework, in which increased firing or synchrony reflects amplified cortical pain processing (Cichon et al., 2017; Eto et al., 2011; Okada et al., 2021). Noxious stimulation reliably engages S1 in humans (Bushnell et al., 1999; Peyron et al., 2000; Ploner et al., 1999). Yet across sensory cortices, activity in awake animals is strongly shaped by movement, arousal, and internal state (Musall et al., 2019; Reimer et al., 2016; Saleem et al., 2013; Stringer et al., 2019). It was unclear how stimulus information is organized relative to movement- and state-related activity within S1 populations during noxious processing, and how injury alters this organization. We measured stimuli, population activity, and behavior on single trials and separately silenced S1 during stimulation. S1 hindlimb activity was dominated by movement and state, stimulus information was carried in latent population dimensions, and inflammatory injury reorganized this structure, making S1 necessary for mechanical hypersensitivity and hyperarousal.

### S1 organizes stimulus and state signals in population dimensions

S1 activity tracked movement and state at both macroscopic and single-neuron levels. Pupil dilation and facial expression predicted neural activity more strongly than movement of the stimulated limb, indicating that S1 activity did not simply reflect local movement or reafference. In barrel cortex, activity carries decision- and outcome-related signals during tactile learning (Bale et al., 2021; Buetfering et al., 2022) and reflects body-state variables such as head and body configuration (Gantar et al., 2025; Palacio-Manzano et al., 2026). During S1 somatosensory processing, movement, arousal, and protective behavior were embedded within the same population activity as stimulus information.

Despite this strong state dependence, population dimensions beyond the state axis carried mechanical stimulus structure. PCA revealed a dominant movement- and state-related axis, while additional dimensions carried stimulus content. Mechanical stimulus content was carried in population dimensions beyond the dominant state-related axis, whereas individual neurons, including those identified by selective optogenetic activation of nociceptors, showed little stimulus selectivity. This is consistent with stimulus information being distributed across low-dimensional population structure rather than expressed as clear single-cell selectivity (Churchland et al., 2012; Elsayed et al., 2016). More broadly, tactile input reaching S1 is already integrated across multiple low-threshold mechanoreceptor classes (Emanuel et al., 2021). In barrel cortex, self-generated tactile reafference from whisking redirects noxious-evoked population trajectories toward non-nocifensive states (Lu et al., 2022). Our analysis addressed a complementary question by isolating mechanical stimulus identity from population dimensions beyond the dominant movement- and state-related axis. Together, these findings identify population organization as an important level at which sensory input is integrated with behavioral state.

### Noxious heat responses in S1 reflect behavioral state rather than stimulus content

Heat responses provided the clearest dissociation between stimulus content and state. The evidence was threefold: (1) heat-evoked macroscopic activity collapsed under anesthesia to a degree comparable to an auditory tone, whereas mechanical and cold responses remained; (2) the population dimensions beyond the state axis carried substantially more stimulus structure for mechanical than for thermal heat input, whether measured as response magnitude, trial-to-trial variability, or separability from conditions without hindpaw stimulation; and (3) S1 inhibition did not reduce heat- or warmth-evoked responses after CFA. Heat therefore evoked arousal- and movement-related activity in S1 hindlimb, but little stimulus-specific structure. This fits evidence that innocuous warmth is represented in posterior insular cortex rather than S1 (Vestergaard et al., 2023), while human scalp-EEG studies show that the large vertex potential evoked by noxious laser heat is shaped by stimulus salience rather than nociceptive specificity (Iannetti et al., 2008; Mouraux and Iannetti, 2009; Ronga et al., 2013). Here, heat still evoked arousal and silencing S1 left this intact. S1 influenced behavior only for mechanical stimuli, whose identity was carried in the population alongside state. S1 therefore shaped the coordinated response only when stimulus information and state were organized within the same population. It did not do so when state-related activity was present with little stimulus information.

This distinction is relevant for cortical pain readouts. S1 calcium signals can classify pain and analgesic states, including in analyses that consider movement (Bak et al., 2024; Yoon et al., 2022), and distributed neuroimaging signatures have motivated a search for brain-based pain biomarkers (Davis et al., 2017; Wager et al., 2013). In S1, classification may rely on combinations of stimulus-, movement-, arousal-, and other state-related signals that covary with pain under the conditions tested. Similar combinations may also occur during fear or other salient states, so successful classification does not reveal which components of S1 activity drive the result. S1 may therefore provide a rich set of signals for decoding pain-related conditions, but interpreting such readouts requires knowing what information they contain.

### Inflammatory injury reorganizes S1 population geometry

Persistent pain has been associated with S1 synaptic remodeling, increased activity, altered inhibition, and synchrony (Cichon et al., 2017; Eto et al., 2011; Kim and Nabekura, 2011; Okada et al., 2021). These findings establish that inflammatory injury changes S1, but do not specify how stimulus- and state-related signals are arranged within the population. After CFA, response magnitude and single-neuron selectivity changed little. Instead, the population eigenspectrum redistributed, participation ratio increased, the state axis became tightly linked to paw movement, stimulus contrast alignment decreased, and co-activity spread across dimensions. Similar changes in spontaneous activity indicated tonic reconfiguration rather than altered evoked gain alone. Reports of S1 hyperactivity, synchrony, or gamma-band pain signals may therefore describe complementary aspects of the same injury-induced cortical state (Yue et al., 2025), consistent with the view that primary sensory cortex is adaptive and context-dependent (Waiblinger et al., 2026). As manifold-based approaches are increasingly used to read out and modulate cortical activity (Busch et al., 2026), clinical translation may depend on which signals are present in S1, how they are organized, and which process a readout or intervention is intended to control.

### S1 sustains coordinated mechanical hypersensitivity and hyperarousal

Optogenetic inhibition showed that during inflammatory injury, S1 is needed for a coordinated mechanical response state. After CFA, inhibition reduced mechanically evoked paw movement, whisking, pupil dilation, and noxious facial expression, while having little effect before CFA and none on heat or warmth responses. Hyperarousal here refers to this coordinated increase in state-related readouts after injury, including autonomic and facial components, rather than to a single hidden state. Earlier work on S1 in chronic pain has centered on paw withdrawal (Cichon et al., 2017; Liu et al., 2018). By measuring several features together, we found that S1 sustains a response extending beyond the paw. S1 carried stimulus and state together before injury, yet inhibition affected these responses after inflammation, suggesting that after injury S1 comes to drive adaptation of widespread protective responses. This could involve descending control of spinal tactile gain, cortico-cortical pathways to affective circuits, or disruption of the population organization that links mechanical input to state (Liu et al., 2018; Singh et al., 2020; Wang et al., 2025). Pain is closely linked to vigilance and defensive behavior, from hypervigilance in humans to injury-induced threat sensitivity in animals (Crombez et al., 2005; Crook et al., 2014; Khuong et al., 2019; Lister et al., 2020; McDermid et al., 1996). S1 inhibition therefore acted less as a selective block of paw output than as a disruption of the inflammatory injury-dependent state coupling mechanical input to hypersensitivity and hyperarousal.

### Limitations

Our recordings targeted S1HL, so we cannot exclude a spatially displaced thermal representation elsewhere in S1 (Osaki et al., 2022), although multiple lines of evidence indicate that heat-evoked activity at the sampled site was dominated by movement and state. Inhibition silenced S1 broadly, so we cannot conclude that geometric reorganization itself is required for hypersensitivity, only that S1 activity is necessary for the coordinated mechanical hypersensitivity and hyperarousal observed after inflammation. Dimension-specific perturbations will be needed to test whether particular axes of stimulus-state embedding are causal.

## Conclusion

These findings distinguish stimulus-state embedding from simple modulation of sensory responses. Movement and arousal did not merely modulate S1 responses to noxious stimuli but also occupied the same population space in which mechanical stimulus identity was represented. At the single-neuron level this appeared as broad state-dependent activity with little reliable stimulus selectivity, whereas at the population level mechanical stimulus identity was carried in coordinated activity across neurons, a dissociation that requires a population-level account (Barack and Krakauer, 2021). The causal results followed the same pattern. S1 inhibition altered the mechanical response, where stimulus and state were embedded together in the same population, but not the response to heat, where state was present without stimulus information.

S1 shaped behavior only where stimulus content and behavioral state were carried together. Inflammatory injury reorganized this embedding into an *injury mode*, in which the state axis and protective movement became more tightly aligned while thermal responses remained state-driven and unaffected by S1 inhibition. S1 therefore does not act simply as a relay that encodes stimulus features and transmits them downstream – it operates within a loop linking the immediate environment, internal state, and protective behavior. Because inflammatory injury changed how activity was organized rather than how much was present, cortical approaches to chronic pain may need to target the population structure that couples mechanical input to state, not only overall S1 excitability.

## Acknowledgements

We are especially grateful to Matteo Carandini for his early advice and support. We thank Mehmet Fişek for discussions on the two-photon recordings, Michael Krumin and Andrew J. Peters for technical advice, and Anyi Liu for initial optogenetic inhibition experiments. Finally, we thank Thomas D. Mrsic-Flogel for donating VGAT-ChR2-EYFP breeder mice.

This work was supported by a Sir Henry Dale Fellowship jointly funded by the Wellcome Trust and the Royal Society (109372/Z/15/Z), and grants from the Royal Society (RSG\R1\180186), the Wellcome Trust (204841/Z/16/Z), and the Medical Research Council (MR/N013867/1).

## Contributions

A.S.P. and L.E.B. conceptualized the study, developed the experimental methodology, curated the data, and performed formal analysis. A.S.P. performed all experiments and wrote the original draft. I.P. trained and validated the DeepLabCut models. L.E.B. built the experimental platform, developed the software, provided resources, supervised and administered the project, and acquired funding. A.S.P. and L.E.B. reviewed and edited the manuscript.

## Methods

### Animals

All animal procedures were approved by the Animal Welfare and Ethical Review Board at University College London and performed under licenses released by the UK Home Office following appropriate ethics review and in accordance with the Animals (Scientific Procedures) Act 1986.

For the widefield calcium imaging experiments, we used transgenic mice expressing GCaMP6s across excitatory neurons by crossing CaMK2a-tTA mice (heterozygous; JAX #007004) and tetO-GCaMP6s mice (heterozygous; JAX #024742). For the optogenetic inhibition experiments, we used transgenic mice expressing channelrhodopsin-2 (ChR2) and EYFP in GABAergic interneurons using VGAT-ChR2-EYFP mice (Zhao et al., 2011) (heterozygous; JAX #014548). For the two-photon calcium imaging experiments, we used transgenic mice that express ChR2 in primary afferent nociceptors, by crossing Trpv1-IRES-Cre mice (Cavanaugh et al., 2011) (heterozygous; JAX #017769) and Cre-dependent ChR2-tdTomato mice (Madisen et al., 2012) (homozygous; JAX #012567). The resultant TRPV1-Cre::ChR2 mice expressed GCaMP6s in primary somatosensory cortex (S1) neurons following stereotaxic injection of AAV1-GCaMP6s (see *Surgery* below).

All experiments used replicates across multiple trials and multiple litters. Mice were given ad libitum access to food and water and were kept at a 12 h light : 12 h dark cycle at 22 ± 2°C and close to 55% relative humidity.

### Surgery

Mice were anesthetized with a mixture of oxygen and isoflurane (2%), injected with buprenorphine hydrochloride (Vetergesic, 0.1 mg/kg), and local anesthesia was provided topically with lidocaine (0.1%) and bupivacaine (0.025%). A clear-skull preparation (Guo et al., 2014) was used for widefield calcium imaging and optogenetic inhibition experiments, and an optical window for two-photon calcium imaging experiments following stereotaxic injection of AAV.

For the clear-skull preparation, used in widefield calcium imaging and optogenetic inhibition experiments, a custom-made U-shaped aluminum headplate was positioned on the edge of the occipital bone and secured to the skull with dental cement (Superbond C&B L-type Radiopaque, Sun Medical). A 3D-printed U-shaped 10 × 10 mm acrylic retainer was attached to the skull and the skin with tissue adhesive (GluTure) to form a boundary anterior to the head-plate. A thin layer of cyanoacrylate glue (Zap-A-Gap) was applied to the skull and allowed to dry. Fast UV-curing optical adhesive (Norland 81) was applied to the skull and cured for 10 min with a 340 nm LED (Thorlabs, M340L4) at 60 mW. Bregma was marked on the optical adhesive. Widefield calcium imaging informed the precise location of S1 activated by hind-paw mechanical stimuli (−0.7 mm AP and 1.85 mm ML relative to Bregma). This location was used to determine the placement of the optical window for two-photon calcium imaging experiments.

For the optical window, a custom-made aluminum headplate with a 7 mm circular aperture was centered on the previously determined S1 coordinates and secured to the skull with dental cement. In the center of the aperture, a 3 mm-diameter craniotomy was created and the exposed brain cleaned with a sterile saline solution. pAAV.Syn.GCaMP6s.WPRE.SV40 was a gift from Douglas Kim and GENIE Project (Addgene, #100843-AAV1, RRID:Addgene_100843 (Chen et al., 2013)) and was diluted 1:8 (0.29 × 10^12^ vg/ml) and injected at 200 nl/min at sites surrounding −0.7 mm AP and 1.85 mm ML (relative to Bregma), 400 µm below the pial surface of the brain.

Cohort 1 received five injections of 200 nl each and cohort 2 received three injections of 300 nl each. The brain surface was covered with an optical window, which comprised 3 mm and 4 mm circular borosilicate glass coverslips sealed together with UV-curing optical adhesive (Norland 61). Dental cement was used to secure the window to the skull. Animals were allowed to recover for at least one week before 1–2 weeks of habituation.

### Set-up for awake behaving head-fixed mice

Mice were head-fixed to a custom frame attached to a manipulator and a stabilization arm that supported the position of the mouse offset from the center of a circular perforated steel-sheet floor (McMaster-Carr) freely rotating on a mounted ball-bearing axis as the mouse walks and moves. Mice were illuminated with three infrared light sources. White noise (70 dB, 44 kHz sample rate) was played to mask background noise. Mice were habituated over 1–2 weeks until they could walk with ease on the rotating grid floor while head-fixed. Each recording session lasted 1–1.5 h.

### High-speed recordings of behavior

Behavior was recorded using two machine-vision cameras (FLIR Blackfly S BFS-U3-04S2M-CS). One captured the face of the mouse using an 8 mm lens (VS-0814H, VS Technology) and IR bandpass filter (BN850-27, MidOpt). The other imaged the body with a 35 mm lens (SV-3518V, VS Technology) and IR bandpass filter (BN850-35.5, MidOpt). These IR bandpass filters reduced reflected laser light that can affect markerless pose estimation. Two four-LED 850 nm light sources illuminated the face and body from each camera, and an additional 15-LED 850 nm light source provided ambient illumination. Each camera frame was triggered through the camera GPIO by an Arduino programmed to generate TTL pulses at 400 frames per second (fps) for 5 s and then 40 fps for 5 s, on receiving a LabVIEW-timed NI multifunction I/O device (NI USB-6343) trigger. Each frame was stored as a time-stamped uncompressed TIFF image for later analysis.

### Laser stimulation module for ST-optogenetics and optical stimulation of skin

A miniaturized laser stimulation module was modified from scanned transdermal optogenetic (ST-opto) control (Parkes et al., 2026; Schorscher-Petcu et al., 2021). This provides precise targeting of laser beams to the hindpaw for optogenetic and infrared thermal stimulation. For transdermal optogenetic stimulation, the beam of a 473 nm laser (Coherent OBIS 473LX) was beam expanded 3× (GBE03-A, Thorlabs) and aligned with two dielectric mirrors (KM100-E02, Thorlabs) to a lens (AC254-500-A, Thorlabs), a 765 nm long-pass dichroic (FF765-DI01- 25X36X2.0, Semrock), an 805 nm short-pass dichroic (DMSP805R, Thorlabs), and two silver galvanometer mirrors (Cambridge Technology 6200H) to a large elliptical dielectric mirror (BBE2- E02, Thorlabs). For infrared thermal stimulation, the beam of a 785 nm laser (SLOC Lasers RLM785TA-1500) was shuttered (SH05/M, Thorlabs), focused (AC508-750-B, Thorlabs) and aligned with two near-IR mirrors (KM100-E03, Thorlabs) to a cylindrical lens (LJ1144RM-B, Thorlabs) and a silver mirror (PF10-03-P01, Thorlabs), sharing the optogenetic stimulation laser beam path to pass through the same 765 nm long-pass dichroic, 805 nm short-pass dichroic, galvanometers, and elliptical mirror. The optogenetic laser beam and infrared laser beam were aligned and focused to give mean spot diameters of 1.25 mm and 4.5 mm, respectively, measured with a beam profiler (BP209, Thorlabs). A putative stimulation site was targeted by ‘descanning’ reflected ambient infrared light back through the galvanometers, reflected at the same 805 nm short-pass dichroic to a camera (FLIR Blackfly S BFS-U3-04S2M-CS) with an 8 mm lens (VS-0814H, VS Technology) streamed in NI LabVIEW. Laser stimulation was time-locked by triggering from the NI multifunction I/O device (NI USB-6343). This approach enables somatosensory stimuli (thermal and optogenetic) to be delivered optically with spatiotemporal precision (Parkes et al., 2026; Schorscher-Petcu et al., 2021).

### Stimulus delivery

All stimuli were delivered in sets, randomly ordered prior to the experiment. Three or more of a stimulus type would be followed by a different stimulus type depending on this pre-assigned random order that differed across sessions for every mouse. Each stimulation was separated by a >2 min interval. A trial was viewed as valid only if the animal was not moving immediately prior to stimulus delivery; this assessment was made online during the recordings and validated in the offline behavioral analysis pipeline.

A diverse range of somatosensory stimuli was delivered to the plantar surface of the hindpaw. For thermal heat stimuli, a 300 ms infrared light pulse was targeted using the 785 nm laser at 4.1 W (‘warmth’, estimated peak 41°C) or 10.1 W (‘heat’, estimated peak 58°C). The laser spot size was 4.5 mm in diameter, and the plantar surface of the hindpaw was darkened with a black marker pen to improve light absorption. Temperatures were estimated using a 1.4 mm × 0.8 mm calibrated black marker-darkened thermistor. For cold stimulation, a drop of acetone was transferred from the tip of a glass rod to the plantar surface of the paw (‘cold’). For mechanical stimuli, the plantar hindpaw was exposed to brief stimulation with a 0.4 g von Frey filament (‘von Frey’); a 30 ms air puff at 50 psi delivered with a pico-spritzer (Picospritzer IID, General Valve Corporation) through a blunt needle positioned around 5 mm from the paw (‘air-puff’); or a blunt insect pin briefly touching the plantar hindpaw (‘pin-prick’). Transdermal optogenetic stimulation of TRPV1::ChR2 primary afferent neurons (‘ST-opto’) was performed using a single-shot 3 ms pulse with a 473 nm laser at 1.25 mm diameter spot size and 40 mW/mm^2^ (Coherent OBIS 473LX), targeted to the plantar surface of the hindpaw. In some experiments, a 300 ms 8 kHz sinusoidal tone at 91 dB (44 kHz sample rate) was delivered as a non-somatosensory stimulus (‘tone’), using an amplified (QTX Q240) NI I/O device analog output to an ultrasound speaker (Pettersson L60). Stimulation controls included ‘off-target’ (a 300 ms infrared laser pulse at 10.1 W aimed proximally but off the hindpaw), ‘approach’ (the hindpaw is approached with a von Frey filament as if to stimulate but without touching the animal), and ‘none’ (trials where stimuli were omitted).

Automatically delivered stimuli (‘heat’, ‘warmth’, ‘ST-opto’, ‘air-puff’, ‘tone’, ‘off-target’) were triggered by the NI multifunction I/O device (NI USB-6343) controlled by custom software written in NI LabVIEW. Manually delivered stimuli (‘cold’, ‘von Frey’, ‘pin-prick’) were post-hoc visually inspected with high-speed videos to determine precise timings. The two behavioral cameras acquired at 400 fps from −3 to +2 s relative to stimulus onset and at 40 fps from +2 to +10 s. To guarantee a consistent 400 fps window across trials for these stimuli, response-window analyses use a conservative cut-off of +1.5 s post-stimulus.

### Widefield calcium imaging

Widefield imaging was performed at 20 fps at 6.2 µm/pixel across 4.3 × 3.2 mm^2^ (696 pixels × 850 pixels) using a Rolera-XR digital CCD camera (Q-Imaging). Hindlimb S1 was targeted by centering the field of view on its approximate location. The area was imaged through a 4× microscope objective (4× Olympus Plan N, NA 0.1, working distance 18.5 mm) on a modified Scientifica microscope focused 400 µm below the pia surface of the brain. Light from an LED (470 nm, CoolLED pE-300white) passed through a filter (ET470/40, Chroma) and long-pass dichroic (T495LP, Chroma) to illuminate the brain. Emitted light passed through the same 495 nm long-pass dichroic, a filter (ET525/50m, Chroma), and 0.5× Olympus tube lens before reaching the camera. Recordings during spontaneous movements did not show aberrant activity in transgenic mice (Steinmetz et al., 2017). Trials were triggered in custom LabVIEW code, which controlled a multifunction I/O device (NI USB-6343) to provide common triggers for the widefield calcium imaging camera and the two high-speed behavioral cameras, enabling synchronization (see *Synchronization* below).

Experiments were conducted on eight mice from three separate cohorts (5 male, 3 female; 11– 21 weeks of age at the first imaging session), recording at least six sessions across six different days for each mouse. In these sessions, we used triplicate trials of multiple stimulus types. For six of the eight mice, the final session was conducted under light sedation and anesthesia using chlorprothixene (5 mg/kg i.p.) and urethane (0.5–1.5 g/kg i.p.), resulting in complete loss of movement for all stimuli used; all stimuli except ‘approach’ were delivered nine times and recordings acquired as before.

### Two-photon calcium imaging

Two-photon calcium imaging was conducted with a modified Scientifica galvo-galvo SliceScope using a 16× water-immersion objective (Nikon CFI LWD Plan Fluorite, NA 0.8, working distance 3 mm). A Ti:Sapphire laser (Coherent Chameleon Ultra II) was used to excite GCaMP6s at 920 nm through a 665 nm long-pass dichroic. Emitted light reflected off the same long-pass dichroic, a second long-pass dichroic (565LP, Chroma), a filter (ET525/50m, Chroma), and two filters (ZET405/473-488/NIRm, Chroma) to block blue masking pulses and blue optogenetic stimuli, before reaching a GaAsP detector (H10770(P)A-40, Hamamatsu). The field of view was 512 × 512 µm (256 × 256 pixels) and images were acquired at 6.1 fps. Deep layer 2/3 of the hindlimb primary somatosensory cortex was targeted by first focusing on the vasculature at the center of the optical window, then descending 320–380 µm below the pial surface (median 355 µm across analyzed sessions). Ambient 1 s blue-light masking flashes were provided at random intervals (mean 10 s) throughout all sessions, except for 12 s from the start of each 10 s trial. Two-photon recordings were acquired continuously throughout experimental sessions. Every two-photon frame and every behavioral camera frame were TTL time-stamped and recorded on a Digidata 1440A (Axon CNS Molecular Devices) to enable synchronization (see *Synchronization*). The two-photon experiments were conducted in two cohorts of mice (2 male, 3 female, imaged at 12–26 weeks of age).

### Optogenetic inhibition

Optogenetic activation of S1 inhibitory interneurons was performed by positioning a 400 µm-core optical fiber (NA 0.39, Thorlabs M82L01) directly above hindlimb S1 in the right hemisphere. This location was the coordinate identified by widefield calcium imaging in the clear-skull preparation where Bregma is visible. A circle of around 5 mm diameter was marked above this location (−0.70 mm AP and 1.85 mm ML relative to Bregma) in the clear-skull preparation to target the optical fiber. Light from a 473 nm laser (Coherent OBIS 473LX) was directed into the back of the optical fiber to provide 1 mm diameter illumination over S1. In optogenetic silencing trials, custom software written in LabVIEW controlled the NI multifunction I/O device (NI USB-6343) to provide a 1 s pulse of light, time-locked to stimulus onset via the same TTL that initiated the sensory stimulus, modulated as a 40 Hz sinusoid with a peak intensity of 1.5 mW at the fiber tip and linearly attenuated in the final 100 ms to avoid post-stimulus rebound firing (Guo et al., 2014). Ambient blue-light flashes were used as above. As an additional spatial control, the optical fiber was moved above the opposite (left) hemisphere while pin-prick was delivered and had no significant effect.

Experiments were conducted on six VGAT-ChR2-EYFP mice across two cohorts (4 female, 2 male; 27–32 weeks of age at the first session). For each mouse, we used one pre-CFA (‘Naive’) session and one post-CFA session (see *Inflammatory injury model*). Within every session, each stimulus was delivered four times: two control trials (no blue-light pulse) and two optogenetic inhibition trials (paired with the 1 s blue-light pulse), interleaved within the stimulus sequence.

### Synchronization

Behavioral recordings, neural recordings, somatosensory stimuli, tones, and masking flashes were synchronized using common triggers. LabVIEW code initiated each trial; the multifunction I/O device (NI USB-6343) directly controlled tones and masking flashes, and drove an Arduino board that was programmed for parallel-port manipulation of each individual frame of each camera (two high-speed cameras for behavior, and one for widefield imaging) at the appropriate frame rate. The same Arduino triggered the somatosensory stimuli at the precise time (3 s after trial onset). In widefield experiments, image-file timestamps were used to validate the timings of neural recordings and account for potential dropped frames, and associated neural and behavioral recordings were paired for each trial. In two-photon experiments, the Digidata 1440A simultaneously recorded two-photon frame TTLs and Arduino TTLs for every individual high-speed camera frame; the ABF stream was parsed with pyabf (Harden, 2022). Trial onsets were identified as the first TTL in each Arduino pulse train separated by more than 30 s from the preceding pulse, and the two-photon frame associated with each trial was the closest frame to stimulus onset +3 s, with ties broken toward the next frame. Any trial manually flagged that the mice were moving during stimulation was excluded from all subsequent analyses.

### Inflammatory injury model

Mice in the optogenetic inhibition and two-photon calcium imaging cohorts underwent an inflammatory injury protocol after the initial 3–6 recording sessions. The day after the last pre-CFA recording, mechanical withdrawal thresholds of the hind limbs were determined with von Frey hairs (up-down method, (Chaplan et al., 1994)). The following day, mice received an injection of 15 µl of 50% Complete Freund’s Adjuvant (CFA) into their left hindpaw. The final recording session took place 24–30 h after CFA injection. All mice showed the characteristic swelling and redness of the injected paw consistent with CFA inflammation.

### Histology

After the final experiment, mice in the two-photon calcium imaging cohort received an intraperitoneal overdose of pentobarbital (Pentoject, 200 mg/kg) and were intracardially perfused with a flush of ice-cold phosphate-buffered saline (PBS) supplemented with 20 IU/ml heparin, then with 35 ml of 4% paraformaldehyde (PFA) in PBS. Brains were obtained and post-fixed in 4% PFA. Brains were sectioned (25 µm, coronal) and imaged by serial two-photon tomography. Image stacks were registered to the Allen Mouse Brain Atlas using brainreg (www.brainglobe.info) and visualized in napari (https://napari.org). The brainseg plugin in napari was used to reconstruct and identify the extent of GCaMP6s expression in S1.

### Behavioral quantification

#### Behavioral video processing and pose estimation

To extract keypoints, high-speed face and body videos were processed offline with DeepLabCut v2.2.0.2 (Mathis et al., 2018) running in an Anaconda virtual environment with Python 3.8.10, TensorFlow-GPU 2.5.0, CUDA 11.2 and cuDNN 8.1. Two markerless pose-estimation networks (ResNet-50) were trained. Videos were selected to represent the full breadth of behavioral responses and *k*-means clustering was used to select the training frames. The face-camera network was trained on 160 manually labelled frames from 8 videos for 200,000 iterations (test-set RMSE 3.19 pixels), with 38 keypoints across pupil, eyelid, mouth, mandibular, nose, ear, whiskers and whisker pad. The body-camera network was trained on 98 manually labelled frames from 4 videos for 200,000 iterations (test-set RMSE 2.91 pixels), with 20 keypoints across tail, forepaws, hindpaws, nose, mouth, ear, and back. Pixel accuracy below 5 pixels on the test set provides good tracking performance comparable to human-level accuracy (Nath et al., 2019). The same two networks were applied to all datasets. Network outputs consist of per-frame *x*, *y* coordinates with a per-keypoint likelihood score.

#### Behavioral metrics

Behavior was quantified from synchronized high-speed face and body videos on a trial-by-trial basis. Metrics fell into two classes. Landmark-based metrics used DeepLabCut tracking and metric-specific likelihood thresholds, where failed frames were set to NaN and linearly interpolated during trace processing. Image-based metrics, including whisker motion and HOG- derived facial-expression traces, were calculated directly from registered video frames and did not use DeepLabCut likelihood thresholds. Unless stated otherwise, traces were aligned to stimulus onset and baseline-subtracted using a pre-stimulus reference window. Scalar response amplitudes were then determined as the mean baseline-subtracted value over the relevant response window. Smoothing was used for display and for threshold-based response detection where specified, but not as a general prerequisite for scalar amplitude extraction.

#### Pupil size

Pupil keypoints on each face-camera frame were accepted when DeepLabCut likelihood >0.5. Frames with the required three key pupil keypoints were fit with an ellipse using the ellipse model implementation in scikit-image. Pupil size was defined as (*a*+*b*)/2, where *a* and *b* are the semi-major and semi-minor ellipse axes. Frames failing the keypoint-likelihood or ellipse-fit criteria were set to NaN and linearly interpolated. The per-trial pupil scalar was the mean baseline-subtracted pupil trace over the relevant response window.

#### Eyelid aperture, blink, orbital tightening and orbital dilation

Eyelid aperture was calculated from three eyelid landmarks when all had DeepLabCut likelihood >0.9. Aperture was defined as the Euclidean distance between upper and lower eyelid landmarks. From the baseline-subtracted aperture trace, three scalar features were derived: blink, defined as the post-stimulus minimum; orbital tightening, defined from negative-going aperture excursions below baseline; and orbital dilation, defined from positive aperture values above baseline. Trials without a positive or negative values for a given component were assigned zero for that component. These features decomposed eyelid dynamics into eye-closure and eye-opening components.

#### Whisker motion

Whisker motion was determined from a rectangular whisker-pad region of interest on the aligned face image. Within this region, frame-to-frame absolute image differences were summed to produce a whisker motion-energy trace. The per-trial whisker scalar was the mean baseline-subtracted motion-energy trace over the relevant response window.

#### Hindpaw kinematics: paw lift and Euclidean paw displacement

The left hindpaw ankle was tracked on the body/stimulus camera and accepted when DeepLabCut likelihood exceeded 0.9. Frames with implausibly large (50-pixel) inter-frame jumps were treated as tracking failures: the flagged frame and the next two frames were set to NaN and linearly interpolated. In the two-photon dataset (Figures 4 and 5), hindpaw points were also filtered frame-by-frame by raw image position: points with *y*-coordinate ≥300 pixels were treated as mistracking and interpolated. This conservative cutoff reduced the chance of accepting points on the visible thermal-laser artifact rather than the paw. In heat and warmth trials, brief thermal-laser artifact frames were also removed and interpolated before paw metrics were determined. Two paw metrics were calculated from the paw trace. Paw lift was the vertical paw displacement. Paw distance was the frame-to-frame Euclidean displacement of the paw in the *x*–*y* plane relative to its pre-stimulus baseline position.

#### Facial-expression scalar traces

Facial-expression metrics were derived from the HOG-based facial analysis pipeline and a random-forest classifier trained on per-trial facial prototypes (see *Face registration and facial expression analysis*). Classifier-derived per-frame probabilities, including noxious-face and tone-face categories, were baseline-subtracted to generate facial-expression traces. The per-trial facial-expression scalar was the mean probability over the relevant response window.

#### Composite behavioral scalars

Two summary measures of overall behavior were derived from the widefield eight-feature matrix (Figure 3): paw lift, paw distance, orbital/eye tightening, orbital/eye dilation, whisker motion, pupil size, noxious face and tone face. Blink was excluded for reasons described under Statistical analysis. ‘Mean behavior’ was the row-wise mean of the *z*-scored behavioral features. PC1_beh_ was the first principal component of the same feature matrix. Both were evaluated as predictors of cortical activity; PC1_beh_ was retained as the primary composite because it gave lower AIC/BIC and higher marginal R2 across models.

#### Analysis windows

In Figures 1−3, behavioral feature matrices used the high-speed peri-stimulus segment over −2,000 to +1,000 ms at 400 fps, unless stated otherwise. For Figures 4 and 5, two-photon population analyses, PC and behavioral scalars used the 0 to +1,500 ms response window where specified, while early neuronal responder analyses used the 0 to +2,000 ms window. For Figure 6 analyses, response scalars were measured over 10–500 ms for rapid responses (paw distance and whisker motion), 10–1,500 ms for slow responses (pupil size), and 10–1,000 ms for mixtures (noxious-face probability). The S1 inhibition baseline window was −1,000 to −10 ms relative to stimulus onset.

### Face registration and facial-expression analysis

Subtle trial-to-trial shifts in the position of the face were corrected by registering each trial to a common reference image. The reference image was a median projection of 500 baseline frames from a representative trial. For each trial, a second median projection was generated from baseline frames −1,500 to −500 ms before stimulus onset. The reference and trial baseline projections were cropped, denoised with a 10 pixel × 10 pixel median filter, and aligned using enhanced correlation coefficient maximization in OpenCV with a Euclidean motion model. Registration used 1,000 iterations and a termination criterion of 10−10, yielding a 2 × 3 affine warp matrix containing translation and rotation parameters. The same transform was then applied to all face-camera frames from that trial.

Registered face images were analyzed using histogram of oriented gradients (HOG), following the general approach of Dolensek et al. (Dolensek et al., 2020). Each aligned frame was cropped to the facial region containing the whisker pad and orbital area. The cropped image stack was temporally smoothed with a Gaussian filter along the time axis (*σ* = 60 frames), and HOG vectors were determined from each frame using 8 orientations, 16 × 16 pixel cells, 1 cell per block and square-root intensity compression. This produced 13 × 29 HOG cells and a 3,016-dimensional HOG vector per frame. The same HOG parameters were used for all datasets.

For each trial, a baseline facial-expression vector was defined as the mean HOG vector over frames −799 to −200 relative to stimulus onset at 400 fps. Post-stimulus HOG vectors from frames 1 to 600 were then ranked by their Pearson correlation with this baseline vector. The 10 post-stimulus frames least correlated with baseline were retained, and their mean HOG vector was used as the per-trial facial-expression prototype. The retained frame indices were stored as the HOG dissimilarity output.

### Random-forest classifier for per-frame facial state

To convert HOG vectors into scalar facial expression traces, random-forest classifiers were trained on per-trial HOG prototypes from the initial awake behavior–widefield imaging experiment used to establish the behavioral and cortical-response relationships in Figures 1–3. This provided an independent facial-expression training set for the later two-photon and optogenetic inhibition. Indeed, facial expression traces in Figures 4–6 were scored by a classifier trained on separate data rather than refit to the experimental condition being tested.

Each training example was one per-trial HOG prototype, labelled by the stimulus identity of that trial. Training labels included heat, pinprick, acetone, von Frey, warmth, air-puff, tone, off-target, approach and no stimulation. Random-forest classifiers used 1,000 trees and were trained repeatedly with different random seeds.

Widefield classifiers were trained on a random 40% of trials within each mouse and tested on the remaining 60%, repeated over 100 random splits. Training used the ten stimulus labels, and predicted labels were grouped post hoc into five categories, giving chance levels that depend on the number of labels contributing to each category (3/10 for noxious and control, 2/10 for innocuous, 1/10 for air-puff and tone). A label-shuffled null was generated by permuting training labels and refitting on each iteration, and categories were compared against this null with Mann– Whitney U tests.

The trained classifiers were then applied frame-by-frame to the HOG vectors from each trial, producing a predicted stimulus-related facial state for every frame. Predicted labels were grouped into broader behavioral categories: noxious face, innocuous face, tone/other face and control face. An air-puff category was also retained for the initial widefield dataset behavioral analyses. The final per-frame trace for each category was the across-repeat mean prediction probability, ranging from 0 to 1 and interpretable as the fraction of classifiers assigning that frame to that facial-state category.

Downstream analyses used these classifier-derived probability traces as facial-expression scalars. The noxious-face trace was used as the main pain-like facial expression measure, and tone/other-face traces were used where specified in widefield behavioral-feature matrices. Per-trial scalar responses were calculated as the mean baseline-subtracted classifier probability over the relevant response window. Label-shuffle controls were generated by refitting classifiers after permuting training labels and repeating the same prediction procedure. These shuffled traces were used as classifier-null references where explicitly reported.

### Analysis of widefield calcium fluorescence imaging

#### Widefield fluorescence processing

Widefield calcium movies were spatially binned 2 × 2, yielding 260 × 260-pixel images at 12.46 µm per pixel. For each trial, image-file timestamps were used to reconstruct the widefield frame-time vector relative to stimulus onset. A baseline fluorescence image was calculated as the mean of the widefield frames falling in the 500 ms before stimulus onset. This timestamp-based procedure accounted for dropped or irregularly timed frames rather than assuming fixed frame indices.

For each mouse, a representative baseline image was selected manually as the registration reference. Each trial baseline image was aligned to this reference by enhanced correlation coefficient maximization in OpenCV, using a translation-only motion model, 1,000 iterations and a termination criterion of 10−10. Contrast-limited adaptive histogram equalization was applied only to the baseline images used to estimate the registration transform, to reduce the influence of illumination differences on intensity-based alignment. The resulting translation warp was then applied to the unenhanced fluorescence frames used for *ΔF/F*_0_ analysis.

Bregma was manually identified in the clear-skull preparation in each mouse’s registration reference image. Per-trial registration translations were combined with per-mouse Bregma coordinates to define the maximum *x* and *y* shifts observed across sessions. These worst-case shifts were used to crop a common 180 × 180-pixel field of view for each mouse, corresponding to approximately 2.24 × 2.24 mm. The crop was anchored so that the relative position of Bregma was matched across animals. In the first cohort, the root-mean-squared registration error between automatic and manual registrations was 4.8 pixels in *x* and 5.0 pixels in *y*, corresponding to an overall two-dimensional error of 6.9 pixels, or approximately 86 µm.

For the *ΔF/F*_0_ calculation, the registered per-trial baseline image was spatially smoothed with a Gaussian filter (*σ* = 30 pixels) to generate *F*_0_. Registered fluorescence frames *F*(t) were then converted pixel-wise to *ΔF/F*_0_ = [*F*(t) − *F*_0_] / *F*_0_. For spatial summary analyses, each 180 × 180 *ΔF/F*_0_ frame was downsampled by 5 × 5 averaging to generate a 36 × 36 cortical activity map. These maps were flattened into 1,296-length vectors when vector-based analyses were required. For trial-level cortical response analyses, a mask-weighted *ΔF/F*_0_ trace was extracted from the S1HL region of interest described below.

For onset and active-site analyses, pixel-wise *ΔF/F*_0_ values were converted to *z*-scores using the mean and standard deviation of the 40 frames, corresponding to 2,000 ms, preceding stimulus onset. Trials were considered to contain detectable cortical activity when more than 100 pixels exceeded *z* > 5 within the first 1,000 ms after stimulus onset. For trials passing this criterion, onset was defined as the first frame within the first 500 ms after stimulus onset in which more than 50 pixels exceeded *z* > 3. Active-site coordinates were extracted from Gaussian-smoothed *ΔF/F*_0_ frames (*σ* = 5 pixels) by taking the *x* and *y* positions of the maxima in the 75^th^-percentile projection profiles. The response peak image was the frame with the largest mean *ΔF/F*_0_ within the first 500 ms after stimulus onset.

#### Cortical regions of interest and quality control

Two circular regions of interest, each with radius 10 pixels, approximately 125 µm, were defined within the registered field of view. A functionally defined ROI was centered on the mean peak response location to hindpaw pinprick across widefield experiments. The S1HL ROI trace was used as the primary cortical activity metric for downstream neural–behavioral analyses. A stereotaxic S1HL ROI was centered on the hindlimb S1 coordinate used throughout the study Trials were excluded if fewer than 20 widefield frames were acquired in the 1,000 ms window after stimulus onset, if the trial was manually flagged because of experimenter error, residual motion or ambiguous stimulus delivery, or if saturation was detected in the scaled 16-bit *ΔF/F*_0_ stack.

To assess field uniformity, mean-intensity profiles were calculated from unaligned no-stimulation trials recorded under anesthesia across six mice. Mean intensity varied by ≤2% across the central field of view along both *x* and *y* axes.

#### Baseline cortex activity-map clustering

For baseline-state analyses, pre-stimulus 36 × 36 cortical activity maps were flattened into 1,296- length vectors and clustered using Ward-linkage hierarchical clustering. Two baseline clusters were defined. Trial fluorescence traces were then sorted by baseline-cluster assignment, and peak response and area under the curve were determined per trace and averaged per mouse.

#### Joint neural–behavioral feature matrix

For trial-level widefield neural–behavioral analyses, behavioral scalars were paired with the S1HL *ΔF/F*_0_ response for each mouse, session, stimulus and trial. Behavioral features comprised paw lift, paw distance, blink, orbital/eye tightening, orbital/eye dilation, whisker motion, pupil size, noxious-face probability and tone-face probability. To assemble this matrix, the neural scalar was calculated from the retained −800 to +399 ms trial window by averaging the S1HL trace from stimulus onset onward, corresponding to 0–399 ms. Features were *z*-scored across retained trials before mixed-effects modeling (see *Statistical analysis*), and trials with missing behavioral or neural values were excluded.

Blink was extracted and included in the behavioral feature set, but excluded from the behavioral PC composite and neural regression models because of collinearity with orbital/eye tightening, as described under Statistical analysis.

#### Multivariate behavioral dispersion

For each stimulus, multivariate behavioral dispersion was calculated from standardized behavioral feature vectors, with one vector per trial. In the analyses relating to Figure 2, these vectors used the nine behavioral scalars: paw lift, paw distance, blink, orbital/eye tightening, orbital/eye dilation, whisker motion, pupil size, noxious-face probability and tone-face probability. Dispersion was defined as the mean pairwise Euclidean distance between all trial vectors for that stimulus.

Two controls were generated. First, a sorted control was obtained by independently sorting each feature column, producing the most coordinated trial structure possible given the marginal distribution of each feature. Second, a shuffled control was generated by independently permuting each feature column, destroying trial-wise feature covariation while preserving each feature’s marginal distribution. Normalized dispersion was calculated as (observed − sorted) / (shuffled mean − sorted). Here, 0 corresponds to the sorted maximum-structure control and 1 corresponds to the independently shuffled null.

#### Behavioral principal component analysis and feature correlations

For widefield fluorescence models, principal component analysis was applied to the eight-feature behavioral matrix comprising paw lift, paw distance, orbital/eye tightening, orbital/eye dilation, whisker motion, pupil size, noxious-face probability and tone-face probability. Blink was excluded. Features had been *z*-scored before PCA. PC1 was retained as the scalar behavioral composite, PC1_beh_, for subsequent mixed-effects regressions. PC1_beh_ was compared with a simple mean behavioral composite in models of cortical activity. The compared models were *activity* ∼ *PC1_beh_* × *Stimuli* + (*1* | *Mouse*) and *activity* ∼ *mean composite* × *Stimuli* + (*1* | *Mouse*), with PC1 retained because it gave lower AIC/BIC and higher marginal *R*^2^.

For analyses relating to Figure 2, a parallel PCA was applied separately within each stimulus using the nine behavioral scalars, including blink, and compared with the sorted and shuffled controls described above.

Feature–feature correlation matrices were calculated separately for each mouse and then averaged across mice. Correlations were calculated as Pearson correlations between trial-wise behavioral scalars, and absolute values were shown as heatmaps. Shuffle controls were generated by independently permuting each feature within mouse and recalculating the same per-mouse correlation matrices.

### Analysis of two-photon calcium fluorescence imaging

#### Two-photon fluorescence preprocessing

Motion correction and ROI detection were performed with Suite2p (Pachitariu et al., 2016). For each session, Suite2p outputs were retained for ROI fluorescence, neuropil fluorescence, deconvolved activity, ROI spatial information and automated cell classification.

Each motion-corrected stack was visually inspected for focal-plane drift, focus jumping and progressive bleaching. Sessions were marked as passed, failed, or usable only up to a manually defined frame. Failed sessions were excluded from further processing, and cropped sessions were analyzed only up to the accepted frame. For the main two-photon analyses, each mouse contributed the matched CFA session and three pre-CFA naive sessions selected from the QC-passed recordings on the basis of imaging stability, absence of focal-plane shifts and usable recording quality. This yielded five mice with three pre-CFA naive sessions and one CFA session per mouse.

For each session, fluorescence was neuropil-corrected as *F*_corr_ = *F* − 0.7*F*_neu_. Corrected fluorescence traces were smoothed with a one-dimensional Gaussian filter (*σ* = 2 frames). ROIs with negatively skewed corrected-fluorescence distributions over the recording were removed, because these traces were dominated by downward fluctuations rather than upward calcium transients.

To reduce the influence of slow fluorescence drift and isolate high-confidence calcium events, we adapted a running-baseline and transient-gating strategy previously used for large-scale two-photon calcium imaging (Low et al., 2014; Zong et al., 2022).

A drift-robust *ΔF/F*_0_ trace was calculated for each remaining ROI using a running-baseline approach adapted from Low et al. and Zong et al. The initial baseline was the centered rolling 8th percentile of *F*_corr_ over a 60 s window. A local standard deviation was calculated over the same window, and low-variance epochs were identified from the rolling 10th percentile of this local standard deviation over 360 s. The mean residual between *F*_corr_ and the rolling percentile baseline during these low-variance epochs was added back to the rolling baseline to generate *F*_0_. *ΔF/F*_0_ was then calculated as (*F*_corr_ − *F*_0_) / *F*_0_.

High-confidence event samples were identified from *ΔF/F*_0_ using a drift-adaptive threshold. For each ROI, baseline mean and standard deviation were estimated from 10 s rolling statistics, smoothed over 60 s and evaluated during low-variance epochs. Event samples were defined as *ΔF/F*_0_ values exceeding mean +4 SD. These event samples were used to gate the Suite2p deconvolved trace, retaining deconvolved activity only at event-like time points. A signal-to-noise ratio was calculated as the mean suprathreshold *ΔF/F*_0_ signal above the 90^th^ percentile of the event distribution divided by the mean absolute first difference of subthreshold *ΔF/F*_0_ samples. ROIs with SNR > 5 were retained and sorted by decreasing SNR.

Three trial-aligned activity tensors were generated. The *ΔF/F*_0_ tensor contained baseline-corrected fluorescence activity. The event tensor contained thresholded *ΔF/F*_0_ events. The main deconvolved-activity tensor contained Suite2p deconvolved activity retained only at time points where the corresponding *ΔF/F*_0_ trace exceeded the adaptive event threshold. Non-event samples were set to zero after trial alignment. This event-gated deconvolved activity tensor was used for single-neuron responder analyses, population PCA, synchrony/co-activity analyses, stimulus decoding and contrast geometry analyses unless stated otherwise.

Trials were aligned to the stimulation frame stored by synchronization. For each trial, 20 s before and 20 s after stimulus onset were extracted, giving 40 s trials of 245 frames at 6.1 Hz. Trial-aligned tensors were generated for eight stimulus classes: heat, pinprick, von Frey, warmth, ST- opto, off-target, approach and no stimulation. Trials with fewer than the expected number of frames were discarded.

#### Two-photon analysis windows

Trial-aligned two-photon tensors covered −20 to +20 s relative to stimulus onset. Analysis windows are reported relative to stimulus onset, unless stated otherwise.

The −10 to 0 s pre-stimulus interval was used for baseline subtraction in signal PCA and stimulus contrast geometry analyses, and as the reference interval for single-neuron responder classification.

For single-neuron analysis relating to Figure 4, the early evoked window was 0 to +2 s. This window was used for early responder classification, response breadth, responder co-occurrence, single-neuron ROC analyses and per-neuron behavioral regression. The late evoked window was +5 to +11 s and was used for late responder classification and early–late activity comparisons.

For population co-activity analyses relating to Figure 5, we used two matched 5 s windows: a baseline window from −10 to −5 s and a peri-stimulus window from −2 to +3 s. These windows were used for synchrony, cosine coherence, PC-removal analyses, signal/noise correlation decomposition and baseline–evoked subspace coupling. Signal PCA for the evoked eigenspectrum used a slightly wider −3 to +3 s fitting window to capture the full stimulus-evoked trajectory.

Scalar stimulus-mean responses, decoder features and PC–behavior correlation features used mean activity over 0 to +1.5 s after stimulus onset. For the scalar PC response heatmap, values were additionally baseline-subtracted relative to −1 to 0 s.

#### Single-neuron responder classification

For each neuron, stimulus and trial, activity was summed over the early evoked window, 0 to +2 s after stimulus onset. This observed evoked sum was compared with a null distribution generated from 1,000 same-duration surrogate windows placed randomly within the −10 to 0 s pre-stimulus reference interval from the same trial. The per-trial *p* value was the fraction of surrogate sums greater than or equal to the observed evoked sum. A neuron was classified as an early responder to a stimulus if at least one trial of that stimulus had *p* < 0.01.

The same procedure was applied to the late evoked window, +5 to +11 s after stimulus onset, to define late responders. Early- and late-responder masks were used for raster ordering, early–late coupling, response breadth, responder co-occurrence, single-neuron ROC discriminability and responder-fraction analyses.

#### Response breadth

For each stimulus, neurons classified as early responders to that stimulus were identified. For each of these neurons, response breadth was defined as the number of stimulus classes, out of eight, for which that neuron was classified as an early responder. Observed breadth values were pooled across the three naive sessions within each mouse, converted to a mouse-level breadth histogram and then averaged across mice.

A matched random-neuron null was generated within session. For each stimulus and session, the same number of neurons as the observed responder cohort was sampled at random from that session. These pseudo-responder neurons were forced to count as responders to the index stimulus, while their real response labels to the other seven stimuli were left unchanged. Breadth was recalculated for the pseudo-cohort over 1,000 random samples. For each mouse, the null histogram used for plotting was taken from the session-level matched null generated by this procedure.

#### Stimulus co-occurrence

Responder co-occurrence was calculated from the early-responder masks. For each ordered stimulus pair A and B, co-occurrence was defined as P(response to B | response to A), the fraction of neurons responsive to A that were also responsive to B. Observed co-occurrence was *z*-scored against the response-breadth-preserving shuffle null. Positive *z*-scores indicate greater-than-expected overlap between responder populations.

#### Single-neuron ROC discriminability

Single-neuron stimulus discriminability was assessed with receiver-operating-characteristic (ROC) analyses using early-window activity as the classifier score. ROC curves and AUC values were calculated for stimulus and modality contrasts and averaged at the mouse level. Label-shuffle controls were generated by permuting stimulus labels within mouse and recalculating AUC.

#### Mean event-activity traces and early-responder proportions

For each stimulus, mean activity traces were averaged across trials and neurons within mouse, then across mice. Additional traces were calculated after restricting to neurons classified as early responders to that stimulus. The fraction of early-responsive neurons was calculated per mouse and stimulus as the number of early responders divided by the number of retained ROIs.

#### Per-neuron behavioral ridge regression

For per-neuron behavioral ridge analyses, Suite2p deconvolved activity was aligned to the behavioral video time base for each trial. Neural activity was extracted around stimulus onset, interpolated onto the 400 fps behavioral frame grid, and restricted to −800 to +399 ms relative to stimulus onset. Paw distance, pupil size and whisker motion traces were baseline-subtracted and aligned over the same window. Neural activity and each behavioral predictor were *z*-scored before fitting.

Ridge regression was fit separately for each neuron and stimulus to model neural activity from three behavioral predictors: paw distance, pupil size and whisker motion. RidgeCV was used for penalty selection over *α* = 0.01, 0.05, 0.1, 0.5 and 1.0 with 11-fold internal cross-validation.

For dominance analysis, neuron × stimulus models with *R*^2^ > 0.15 were included. Each model was assigned to the behavioral predictor with the largest absolute standardized coefficient.

Dominance fractions were calculated within each mouse, condition and stimulus as the number of above-threshold models dominated by each feature divided by the total number of above-threshold models for that mouse, condition and stimulus.

#### Signal PCA and population eigenspectrum

Signal PCA was performed per session on trial-averaged, baseline-subtracted activity. For each stimulus, trials were baseline-subtracted using the mean activity over −10 to 0 s and then averaged. The resulting stimulus-averaged responses were concatenated across stimuli. Each neuron was *z*-scored using its mean and standard deviation over the −3 to +3 s evoked fitting window, and full-rank PCA was fit to the concatenated evoked data. No smoothing was applied before PCA.

Effective dimensionality was quantified from the full eigenspectrum using the participation ratio, *PR* = (*Σλ*_i_)^2^ / *Σλ*_i_^2^, and the fractional number of PCs required to explain 80% of variance. Tail participation ratio was calculated after excluding PC1. PC1–PC3 traces were obtained by projecting trial-averaged activity onto the first three PCs, with signs fixed so that the mean response over 0 to +2 s was positive.

For the scalar PC response heatmap, PC responses were measured as the mean projected response over 0 to +1.5 s after stimulus onset, relative to the −1 to 0 s baseline.

#### Population PC–behavior correlations

Single trials were projected into the signal-PCA space fit for that session. For each trial, PC scores and behavioral scalars were averaged over 0 to +1.5 s after stimulus onset. Behavioral scalars were paw lift, paw distance, pupil size, whisker motion and noxious-face score.

Two correlation analyses were performed. First, pooled PC–behavior correlations were calculated across all trials within session, preserving both stimulus-driven differences and trial-to-trial covariation. Second, stimulus-controlled correlations were calculated by demeaning both the PC score and the behavioral scalar within each stimulus type before pooling trials and calculated Pearson correlations. This removes correlations driven only by differences in mean response across stimulus classes.

Session-level correlations were Fisher-Z transformed before mouse-level averaging and statistical testing. For the naive condition, the three pre-CFA sessions were averaged within mouse. The CFA session was analyzed separately.

#### Spatial structure of PC weights

PC1 and PC2 loading vectors from the signal PCA were assigned to the corresponding Suite2p ROI centroids. Spatial autocorrelation of PC weights was quantified with Moran’s I at radii of 10, 15 and 25 pixels, corresponding to approximately 20, 30 and 50 µm. Moran’s I values were *z*- scored against 1,000 nulls generated by shuffling PC weights across ROI positions.

A complementary same-sign nearest-neighbor statistic tested whether positive- and negative-loading ROIs were spatially segregated. For each component, the mean nearest-neighbor distance was calculated separately among positive-weight and negative-weight ROIs and averaged. This observed distance was compared with a null generated by shuffling weight signs across ROI positions. Negative Z values indicate that ROIs with the same weight sign are closer than expected by chance. Bivariate PC1/PC2 loading distributions were also plotted across ROIs.

#### Population co-activity and PC-removal controls

Population co-activity was quantified on single trials in two matched 5 s windows: a baseline window from −10 to −5 s and a peri-stimulus from −2 to +3 s. Two complementary pairwise measures were used. Pearson synchrony was the mean off-diagonal Pearson correlation between simultaneously recorded neurons after subtracting each neuron’s within-trial mean. Cosine coherence was the mean off-diagonal cosine similarity between neuronal time series after causal exponential smoothing with *tau* = 0.33 s. Smoothing was used for cosine coherence because sparse deconvolved activity can make cosine similarity sensitive to coincident zeros.

For PC-removal controls, a session-level PCA was fit to *z*-scored activity concatenated across trials and stimuli within each analysis window. The specified PCs were projected out of each trial, and Pearson synchrony and cosine coherence were recalculated. PC-removal variants were raw activity, no PC1, no PC2, no PC3, no PC1+2 and no PC1+3. For each component, dPCk was defined as raw co-activity minus co-activity after removing PCk. The PC1 vs PC2 balance metric was defined as dPC1 − dPC2.

For each session and window, neuron time series were independently permuted within trial 100 times to generate a time-shuffle null.

#### Baseline–evoked subspace coupling

To test whether CFA changed the relationship between pre-stimulus population activity and evoked activity, PCA was fit to the −10 to −5 s baseline window for each session. The top 10 baseline-window PCs defined the baseline subspace. Evoked activity from −2 to +3 s was projected into this subspace and its orthogonal complement.

Four metrics were calculated per session: the fraction of evoked variance captured by the baseline subspace, the complementary fraction in the baseline-null subspace, the cosine similarity between baseline-window PC1 and evoked-window PC1, and the Pearson correlation between trial-level projection scores onto baseline PC1 and evoked PC1.

#### Continuous and spontaneous eigenspectrum

For continuous-trace eigenspectrum analyses, the same ROI filtering and SNR-ordering pipeline was rerun from the Suite2p outputs, but the continuous unthresholded Suite2p deconvolved activity was used rather than the trial-aligned event-gated tensor. Neuron counts were checked against the trial-aligned tensor after filtering.

For each session, the continuous deconvolved trace was cropped to the usable recording duration. Spontaneous frames were defined as frames more than 30 s from any stimulus onset. Continuous activity was *z*-scored per neuron using full-trace statistics. PCA with up to 50 components was fit separately to the full trace and to spontaneous frames. Participation ratio, PC1 explained variance, cumulative explained variance and tail participation ratio were calculated. Spontaneous frames were also split into early and late halves as a drift check.

#### Signal and noise correlation decomposition

Signal and noise correlations were calculated in the −2 to +3 s peri-stimulus window for the five somatosensory stimuli: heat, pinprick, von Frey, warmth and ST-opto. Off-target and approach controls were excluded.

Signal correlation measured similarity in stimulus-evoked response patterns. For each neuron, trial-averaged responses were concatenated across the five stimuli, and Pearson correlations were calculated between neuron pairs. The session value was the mean pairwise correlation.

Noise correlation measured shared trial-to-trial residual activity after removing the mean stimulus response. For each stimulus and trial, the residual response was calculated by subtracting the mean of the other trials of the same stimulus. Pearson correlations were then calculated between neuron pairs and averaged across trials and stimuli.

#### Population stimulus contrast geometry

Population stimulus contrast geometry was calculated from the five stimuli: heat, pinprick, von Frey, warmth and ST-opto. Trial counts were balanced within each session by truncating all stimuli to the minimum trial count available for that session.

For each stimulus, a stimulus-mean response vector was calculated as the mean baseline-subtracted response over 0 to +1.5 s after stimulus onset. Baseline was the mean activity over −10 to 0 s. Stimulus-mean response vectors were arranged in a neurons × stimuli matrix. For each neuron, values were *z*-scored across the five stimuli using that neuron’s mean and standard deviation across stimuli. Single-trial residuals were calculated as each trial response minus the corresponding stimulus-mean response vector.

Signal dimensionality was calculated as the participation ratio of the stimulus-mean response matrix singular-value eigenspectrum. Response vector alignments were then calculated for every pair of stimuli. For each pair, the direction vector was the difference between the two *z*-scored stimulus-mean response vectors in *N*-dimensional neuron space. This vector was L2-normalized by dividing each neuron’s weight by the Euclidean norm of the full vector, giving a unit-length stimulus contrast vector. This normalization removes differences in response magnitude, so the analysis tests the angle between stimulus contrasts rather than how far apart the mean responses are. L2 normalization and inner products were implemented with NumPy matrix norm and multiplication.

Response vector alignment was calculated as the mean absolute cosine between all distinct pairs of unit stimulus contrast vector. A value near 1 means that different stimulus contrasts are aligned along similar population axes. A value near 0 means that contrasts are aligned along more orthogonal axes.

Response-vector spread was calculated separately as the mean pairwise Euclidean distance between *z*-scored stimulus-mean response vectors, providing a magnitude-sensitive measure of overall stimulus separation. Noise–signal overlap was calculated as the fraction of single-trial residual variance projecting onto the top three left singular vectors of the stimulus-mean response matrix, providing a measure of how strongly trial-to-trial noise lay within the stimulus-coding subspace.

#### Stimulus structure beyond the state axis

To ask what the non-leading population dimensions carried, two measures were calculated per mouse from the scalar PC scores over 0 to +1.5 s. Response magnitude was the mean absolute PC2–PC3 score across trials of a given stimulus. Trial-to-trial spread was the standard deviation of PC2–PC3 scores across trials of that stimulus. Stimuli were grouped into mechanical (pin- prick, von Frey) and thermal (heat, warmth), averaged within group for each mouse, and compared across mice with an exact paired sign-flip permutation test.

Separation from control conditions was assessed with shrinkage linear discriminant analysis (least-squares solver, shrinkage = 0.5) using leave-one-out cross-validation, decoding trials of a given stimulus from off-target and approach trials. Performance was summarized as balanced accuracy (chance 0.5) and averaged within modality for each mouse.

#### Session-order control

Because the CFA session was always acquired after the three naive sessions, a session-order control was performed on evoked Pearson synchrony. For each mouse, the adjacent naive- session change was calculated as Naive_3_ − Naive_2_, and the CFA transition was calculated as CFA − Naive_3_. The step-excess statistic was defined as (CFA – Naive_3_) − (Naive_3_ − Naive_2_). Step-excess values were tested across mice with an exact sign-flip test over 2^5^ = 32 permutations.

### Statistical analysis

#### Shared conventions

The mouse was always the unit of biological replication. For primary permutation-based analyses, trial- or session-level measurements were first reduced to one value per mouse and condition unless stated otherwise. Confirmatory mixed-effects models were fit to trial-level data with mouse included as a random effect. Sample sizes are reported as the number of mice contributing to each panel.

Primary comparisons were tested with exact permutation procedures rather than parametric tests. Parametric mixed-effects models were used as confirmatory analyses for the S1 inhibition dataset.

#### Exact sign-flip permutation tests

Paired within-mouse comparisons were tested with exact sign-flip permutation tests. For each mouse, the paired difference was calculated. The observed statistic was the mean paired difference across mice. The null distribution was generated by enumerating all 2^n^ possible sign assignments of the per-mouse differences, where *n* is the number of mice. One-tailed or two-tailed *p* values were then calculated from this exact null distribution according to the directional hypothesis specified for each analysis.

For interaction contrasts, the paired difference was first calculated separately for each component contrast within mouse. The interaction value was then the within-mouse difference between these paired contrasts, and this interaction value was tested with the same exact sign-flip procedure.

#### Widefield mixed-effects models and ridge regression

Widefield mixed-effects models were fit in R using lme4::lmer with REML. *P* values were obtained with lmerTest using Satterthwaite’s approximation. Marginal and conditional *R*^2^ were obtained with MuMIn::r.squaredGLMM. The stimulus factor included pinprick, heat, acetone, von Frey, warmth, air-puff, tone, off-target, approach and no stimulation, with no stimulation as the reference level. Benjamini–Hochberg FDR correction was applied within each feature or model family as indicated. Four widefield model families were used.

1. **Stimulus–behavior models.** For Figure 2, each behavioral feature was modeled separately as:

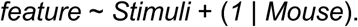 Stimulus-level *p* values were FDR-corrected within each feature.
2. **Stimulus-specific neural models**. For Figure 3 and Supplementary Figure 9, each non-baseline stimulus was compared with no stimulation using:

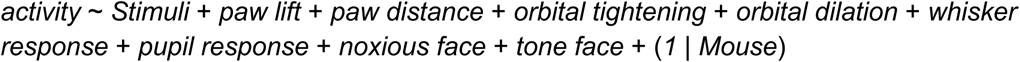 Blink was excluded from neural models because of collinearity with orbital tightening. Variance inflation factors were calculated from the fixed-effects design matrix. If a mixed model was singular, the same fixed-effects model was fit as an ordinary linear model.
3. **Stimulus-agnostic neural models**. Across all trials and stimuli, S1 activity was modeled from behavioral predictors using:

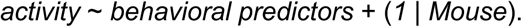 **Behavioral-composite neural models**. The behavioral PC1 composite was tested using:

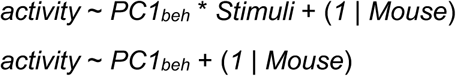 A corresponding total-behavior model was fit in parallel. PC1_beh_ was retained as the primary behavioral composite because it gave better model fit by AIC/BIC and higher marginal *R*^2^. Widefield experiment ridge regression was fit per stimulus in R using glmnet::cv.glmnet with *α* = 0 and standardization enabled. Design matrices contained the behavioral predictors used in the neural models, excluding blink. The regularization penalty was selected as lambda.min from 10- fold cross-validation. Ridge coefficients are reported descriptively. No *p* values were assigned to ridge coefficients.

#### Two-photon exact CFA-relabeling tests

For the main two-photon analyses, each mouse contributed three pre-CFA naive sessions and one CFA session. Exact within-mouse CFA relabeling was used as the primary test. Under the null hypothesis, the CFA label is exchangeable across the four sessions within each mouse. With five mice, this gives 4^5^ = 1,024 exact relabelings.

For univariate session-level metrics, the observed statistic was the across-mouse mean of CFA − mean(naive).

For each null relabeling, one session per mouse was assigned as pseudo-CFA and the remaining sessions were assigned as pseudo-naive. The same statistic was recalculated for all 1,024 relabelings. One-tailed or two-tailed *p* values were calculated from this exact null distribution according to the directional hypothesis specified for each analysis.

For PC-removal co-activity profiles, the relabeling test was applied to nested ordinary-least-squares (OLS) models with mouse as a blocking factor. The reduced model contained mouse and PC-removal terms. The condition model added condition, and the interaction model added the condition × PC-removal interaction. *F* statistics for the condition main effect and condition × PC-removal interaction were compared with the exact within-mouse relabeling null. Because *F* statistics are non-negative, significance was assessed as the fraction of relabeled datasets with an *F* statistic greater than or equal to the observed *F* statistic.

Leave-one-mouse-out robustness was assessed by repeating the exact CFA-relabeling test after dropping each mouse in turn. For co-activity analyses, within-trial time-shuffle nulls were used as a session-level sanity check that observed Pearson synchrony and cosine coherence exceeded values expected after disrupting temporal alignment.

#### Optogenetic inhibition experiment statistical analyses

For Figure 6, the primary test was an exact sign-flip permutation test across six mice. Four mouse-level contrasts were tested for each stimulus × behavior cell: CFA sensitization, inhibition effect before CFA, inhibition effect after CFA and the condition × inhibition interaction. Each contrast was tested one-tailed in the predicted positive direction defined by the contrast sign convention. With six mice, each test used 2^6^ = 64 exact sign assignments.

Confirmatory mixed-effects models were fit in R. Trial-level behavioral data were modeled separately for each stimulus and behavioral feature using:

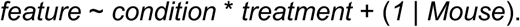

The condition factor contained naive pre-CFA and CFA levels, and the treatment factor contained control and inhibition trials. Type-III F tests were obtained with lmerTest using Satterthwaite’s approximation. Within-condition inhibition effects were estimated with emmeans using pairwise treatment contrasts without multiple-comparison adjustment. Model singularity was recorded for each fit.

Paw response probability was analyzed with a separate binomial GLMM for key stimuli:

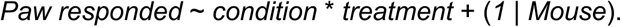

Type-III *χ*^2^ tests were obtained with car::Anova, and within-condition treatment contrasts were estimated with emmeans on the response scale. *P* values from inhibition LMM and GLMM analyses were reported unadjusted.

### Software versions

DeepLabCut training was performed in a dedicated Anaconda environment with Python 3.8.10 on Windows 10 (AMD Ryzen 5 3600 CPU, NVIDIA GeForce RTX 2080 Ti GPU, 64 GB RAM) using DeepLabCut v2.2.0.2 with TensorFlow-GPU v2.5.0, CUDA v11.2 and cuDNN v8.1.

Pre-processing of neural activity and behavior used Python v3.7.11 with numpy v1.21.5, pandas v1.3.5, scipy v1.7.3, scikit-learn v1.0.2, scikit-image v0.19.2, opencv-python v4.5.5.64, tifffile v2021.11.2, imageio v2.18.0, seaborn v0.12.1, matplotlib v3.5.1, and statsmodels v0.13.2. This environment was used for all widefield frame pre-processing, 2P session processing, Suite2p handling, behavioral-video extraction, and HOG analysis.

Analysis (regressions, PCA, decoding, plotting) used Python v3.8.16 with numpy v1.24.2, pandas v1.5.3, scipy v1.10.0, scikit-learn v1.2.2, matplotlib v3.7.0, seaborn v0.12.2, opencv-python v4.9.0.80, statsmodels v0.14.0, and pyabf v2.3.8.

Mixed-effects modeling and ridge regression used R v4.2.3 with lme4 v1.1-35.1, lmerTest v3.1-3, broom.mixed v0.2.9.6, MuMIn v1.48.4, car v3.1-3, glmnet v4.1-8, emmeans v1.10.5, dplyr v1.1.4, and ggplot2 v3.5.0.

### Cohorts and inclusion

Figures 1−3. The widefield experiments comprised eight mice across three cohorts, as described under Animals. All eight mice contributed awake behavioral and widefield data. Six of the eight mice also contributed an anesthesia session (WF07, WF11, WF12, WF14, WF15 and WF16); WF9 and WF13 contributed awake sessions only. Paired awake–anesthesia analyses were therefore restricted to the six mice with both conditions.

Figures 4 and 5. The two-photon experiments comprised five mice across two cohorts: 2P07, 2P08, 2P09, 2P10 and 2P11. All two-photon sessions were first inspected for imaging stability as described under Two-photon preprocessing. For the main naive–CFA analyses, each mouse contributed the matched CFA session and three high-quality pre-CFA naive sessions. The session set used for the main analyses was: 2P07 (Session 3, Session 4, Session 5, CFA); 2P08 (Session 2, Session 5, Session 6, CFA); 2P09 (Session 2, Session 3, Session 5, CFA); 2P10 (Session 1, Session 2, Session 5, CFA); and 2P11 (Session 4, Session 5, Session 6, CFA).

This gave 15 naive sessions and 5 CFA sessions for the two-photon analyses. Mice 2P01–2P06 were pilot animals and were not included in the analyses. Mice 2P05 and 2P06 were used in pipeline-validation.

Figure 6. The optogenetic inhibition experiments comprised six VGAT-ChR2-EYFP mice across two cohorts, as described under Animals. Each mouse contributed one pre-CFA naive session and one post-CFA session. Within each session, control and S1-inhibition trials were interleaved for each stimulus.

## Notes

### Competing Interest Statement

The authors have declared no competing interest.

