## Supplementary Figures for "A stimulus–state geometry in somatosensory cortex reorganizes during inflammatory pain"

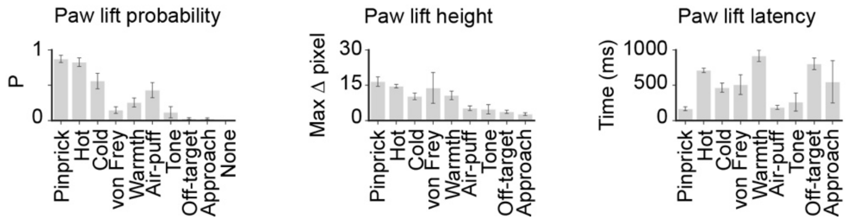


**Supplementary Figure 1 (relates to Fig. 2A). Paw-lift response characteristics across multi-modal stimuli.** Bar plots summarizing the paw-lift response across stimuli. *Left*, paw-lift probability. The per-trial fraction of trials in which the smoothed baseline-subtracted paw-lift trace crossed a threshold during the 0 to +1 s post-stimulus window, averaged per mouse, then across mice. *Middle*, paw-lift height. The maximum $\Delta y$ distance in pixels over the 0 to +1 s response window, per mouse then averaged across mice. *Right*, paw-lift latency. The first frame at which the baseline-subtracted paw-lift trace crossed the response threshold, expressed in ms (captured at 400 fps). Only trials with a detected response contributed. All mean ± SEM (n = 8 mice).

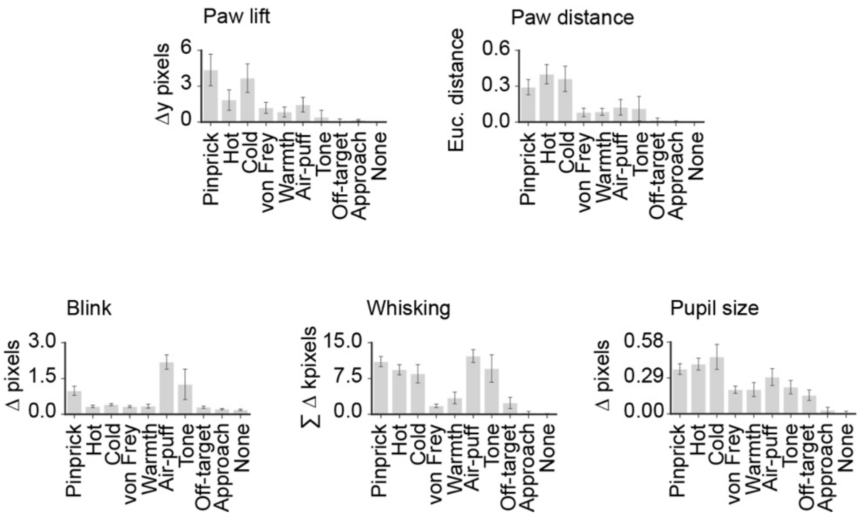


**Supplementary Figure 2 (relates to Fig. 2A). Scalar response magnitudes across different behavioral features.** Bar plots summarizing the per-stimulus scalar responses. Each scalar is the mean of the baseline-subtracted response trace over 0 to +1 s post-stimulus. All mean ± SEM (n = 8 mice).

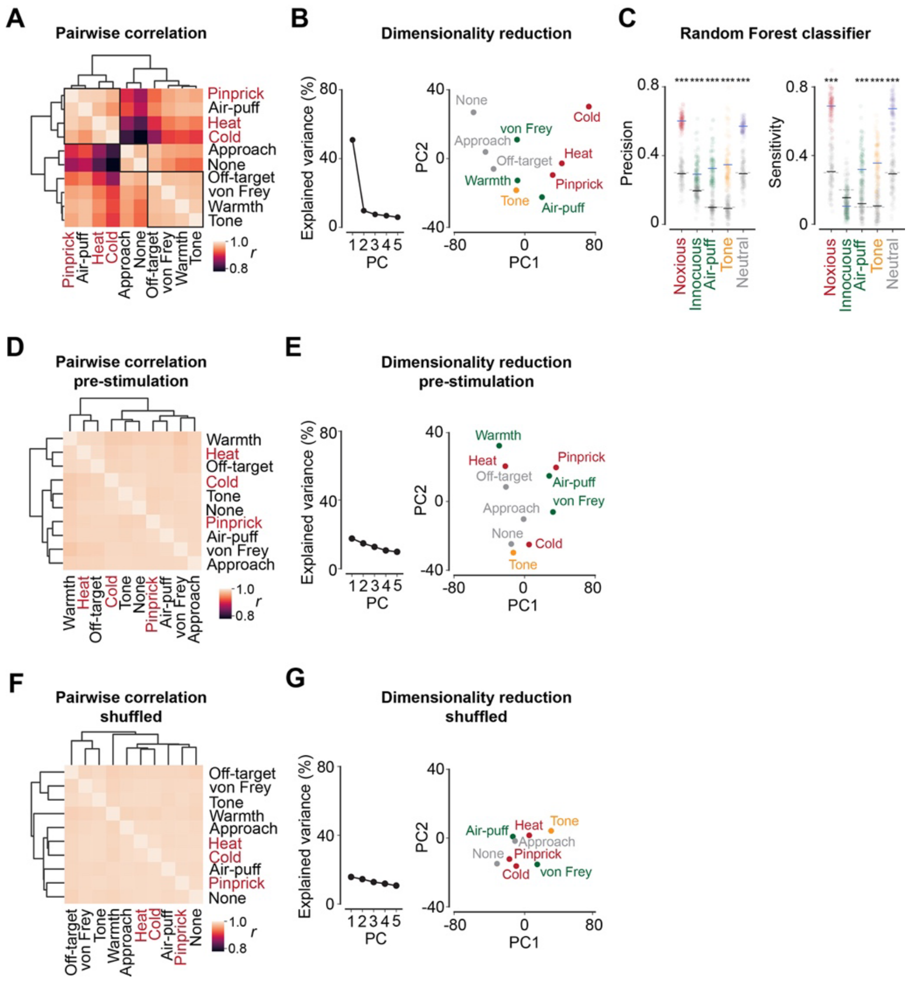

**Supplementary Figure 3 (relates to Fig. 2A). HOG-based facial expression analysis and random forest classifier. (A)** Pairwise Pearson correlation matrix of per-trial prototypical HOG vectors across multi-modal stimuli, pooled across all trials and mice. Dendrograms: Ward-linkage hierarchical clustering on correlation distance. Even using pairwise correlations the noxious stimuli (pinprick, heat, cold in *red text*) clustered together. **(B)** Variance explained and 2D PCA embedding of per-trial prototypical HOG vectors, pooled across trials and mice. *Left*, variance explained by the first five PCs. *Right*, mean per-stimulus position in PC1–PC2 space; dot color by stimulus class. PC1 separated noxious (pinprick, heat, cold) from non-noxious stimuli and controls. **(C)** Random-forest classifier precision for each stimulus category. Classifiers (scikit-learn RandomForestClassifier, 1,000 trees) were trained on per-trial HOG prototypes using a 40/60 train/test split of trials within each mouse, repeated over 100 random splits. Classifiers were trained on ten stimulus labels, and predictions were grouped post hoc into five categories (noxious, innocuous, air-puff, tone, neutral), giving category-dependent chance levels (dashed lines). Dots show individual fits and horizontal lines show mean ± SD across the 100 fits. Grey points show a label-shuffled null generated by permuting training labels and refitting on each iteration. Asterisks mark categories with precision significantly above the shuffled null (Mann–Whitney U). **(D)** Pairwise Pearson correlation matrix of pre-stimulation HOG vectors (500–1500 ms before stimulus onset) across stimuli. Pre-stimulation HOG vectors were weakly correlated across all stimuli, indicating that the inter-stimulus structure seen in (A) emerges only after stimulation. **(E)** Variance explained and PC1–PC2 embedding of pre-stimulation HOG vectors, as in (B). PC1 scores are substantially smaller and mean position structure is absent, consistent with (D). **(F)** Pairwise Pearson correlation matrix of shuffled-label HOG vectors across stimuli, produced by permuting stimulus labels across trials before averaging. **(G)** Variance explained and PC1–PC2 embedding of shuffled-label HOG vectors, as in (B). As expected for the null, structure in panels (A, B) disappeared when labels are shuffled.

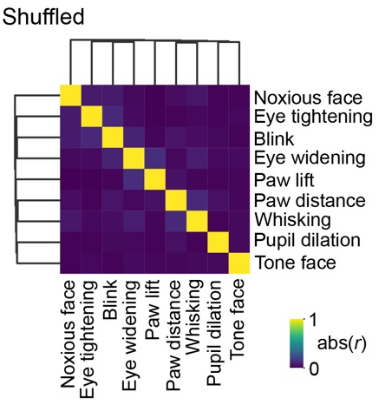


**Supplementary Figure 4 (relates to Fig. 2B). Shuffled-label behavioral correlation null.** Pairwise absolute Pearson *r* matrix of the behavioral scalars calculated after independently permuting the values of each feature across trials within mouse (1,000 shuffles, averaged). Dendrogram: Ward-linkage hierarchical clustering on correlation distance. Off-diagonal values are homogeneously low and the clustering structure seen in Fig. 2B is absent, confirming that the Fig. 2B structure was a property of trial-wise co-variation, not of marginal feature distributions.

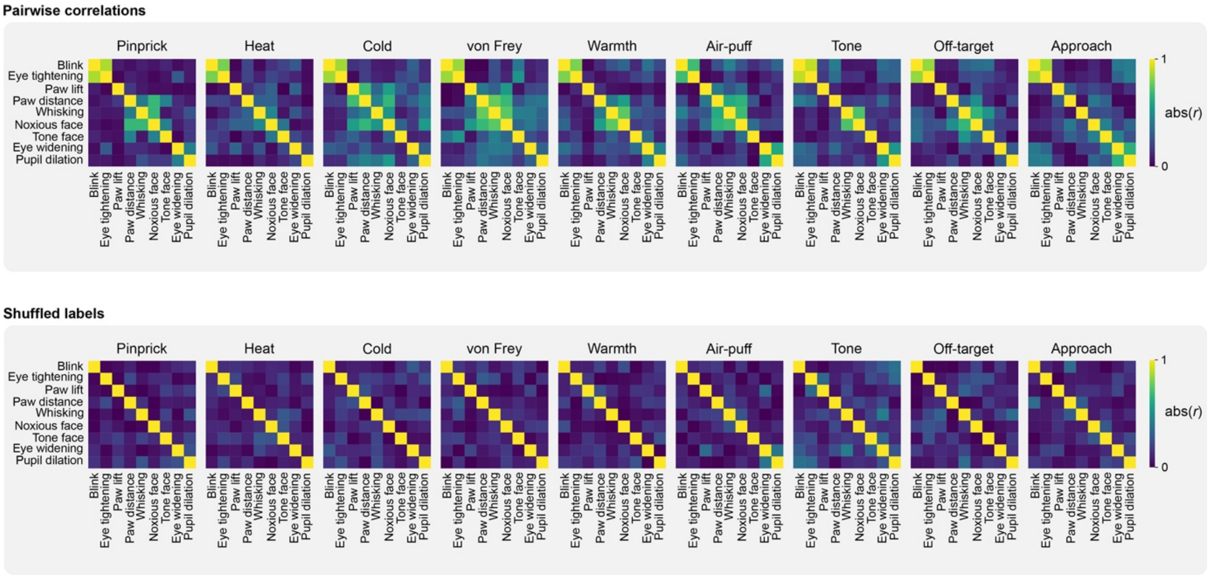


**Supplementary Figure 5 (relates to Fig. 2B). Per-stimulus behavioral pairwise correlations and shuffle null.** Per-stimulus pairwise absolute Pearson *r* matrices of the behavioral scalars. *Top*, observed correlations. Matrices per mouse then averaged across mice for each of the stimuli. *Bottom*, shuffled-label null. Matrices determined after independently permuting each feature within mouse (1,000 shuffles). Stimulus-dependent co-variation structure was clearly visible in the observed matrices (*top*) and absent in the shuffled controls (*bottom*).

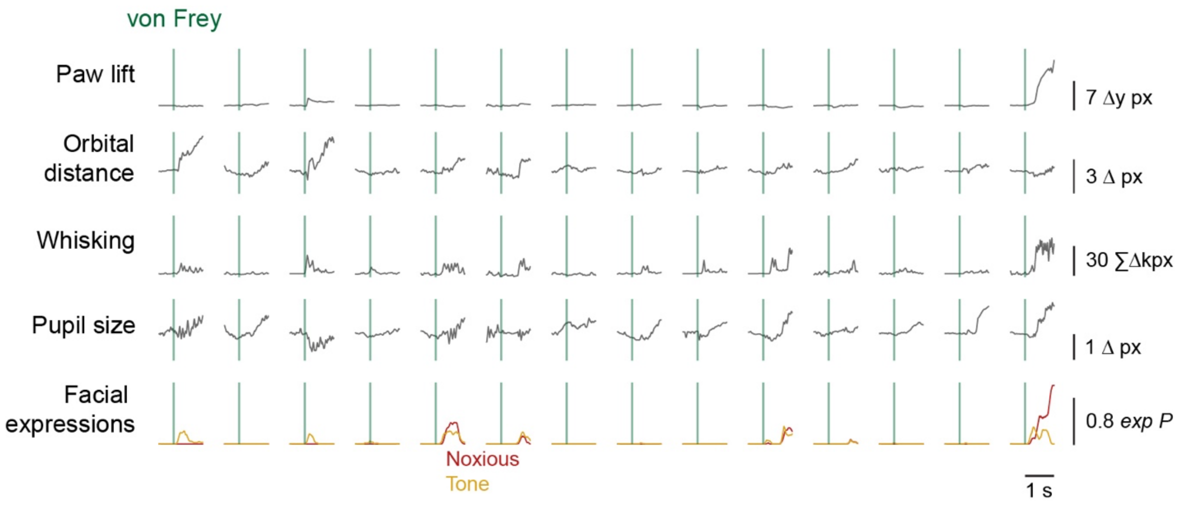


**Supplementary Figure 6 (relates to Fig. 2D). Representative von Frey trials across five behavioral features.** Individual-trial traces for 10 representative von Frey trials (one per column), across five behavioral features. Vertical *green* line, stimulus onset. Across trials, von Frey recruited a highly variable combination of features. Some trials showed only a brief paw movement, others showed pupil size and whisking without paw lift, and others showed strong noxious-face response without a paw response. This was consistent with the high dispersion in Fig. 2F.

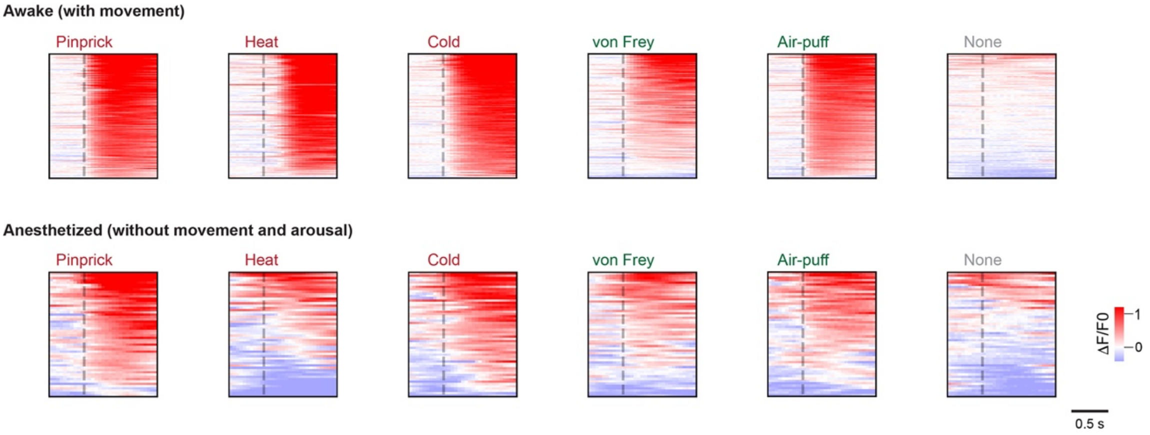


**Supplementary Figure 7 (relates to Fig. 3C). Δ*F*/*F*₀ heatmap time-courses across all stimulus trials.** $\Delta F/F_0$ time-courses for every trial in Fig. 3C averages, shown as heatmaps (*rows* = trials, *columns* = time relative to stimulus onset). *Top*, awake (with movement; *n* = 6 mice, trials pooled); *Bottom*, anesthetized (without movement; *n* = 6 mice). *Dashed* vertical lines are the stimulation onsets. Under anesthesia, responses to thermal heat stimuli dropped while pinprick responses were preserved.

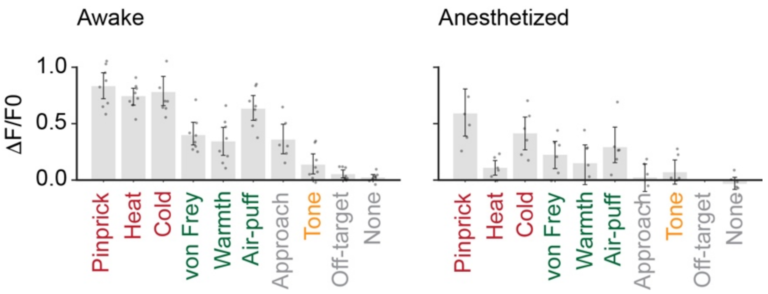


**Supplementary Figure 8 (relates to Fig. 3D). Per-stimulus cortical response magnitude, awake and anesthetized.** Bar plots of mean $\Delta F/F_0$ within S1HL ROI across the 0–500 ms post-stimulus window, per stimulus and condition. Under anesthesia, pinprick, von Frey, air-puff, and cold responses were largely preserved (~70% of awake magnitude) while heat, warmth, tone and approach responses were strongly attenuated. The matched drop across innocuous and auditory stimuli is consistent with the interpretation that awake S1 activation in these classes is movement- and arousal-driven. All show mean ± 95% bootstrap CI ($n$ = 6 mice) where dots are individual mice.

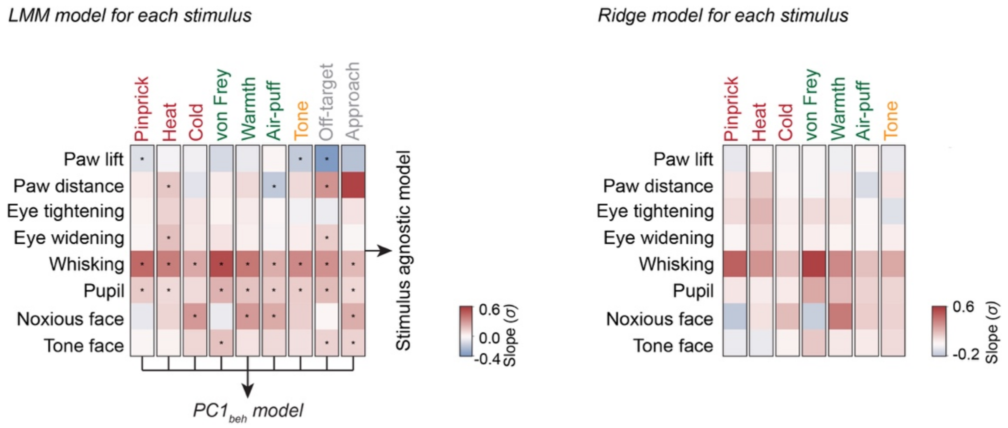


**Supplementary Figure 9 (relates to Fig. 3H). Multi-modal per-stimulus LMM and ridge coefficient matrices.** *Left*, per-stimulus linear mixed-effects model coefficients (σ-units) modeled using *activity* ~ *Stimuli* + *paw lift* + *paw distance* + *orbital tightening* + *orbital dilation* + *whisker response* + *pupil response* + *noxious face* + *tone face* + (*1* | *Mouse*) and fit separately on the two-level subset *{stimulus, no stimulus}* for each of non-baseline stimuli. Arrows indicate the stimulus-agnostic model fit across all stimuli pooled shown in Fig. 3H and $PC1_{beh}$ model in Fig. 3I,J. Asterisks mark BH-FDR-corrected $p < 0.05$ (adjusted within stimulus). *Right*, corresponding ridge-regression coefficients from glmnet 10-fold cross-validation (alpha = 0) at min.lambda. Ridge coefficients were standardized and confirmed the LMM structure: whisking, pupil size, and noxious face dominated across stimuli. No *p*-values are associated with the ridge model. Note: Coefficients reflected relative associations within each stimulus type. Models were fit separately per stimulus, so slopes were not directly comparable across stimuli. Differences in apparent effect size may be influenced by variability or signal strength within each stimulus in this approach.

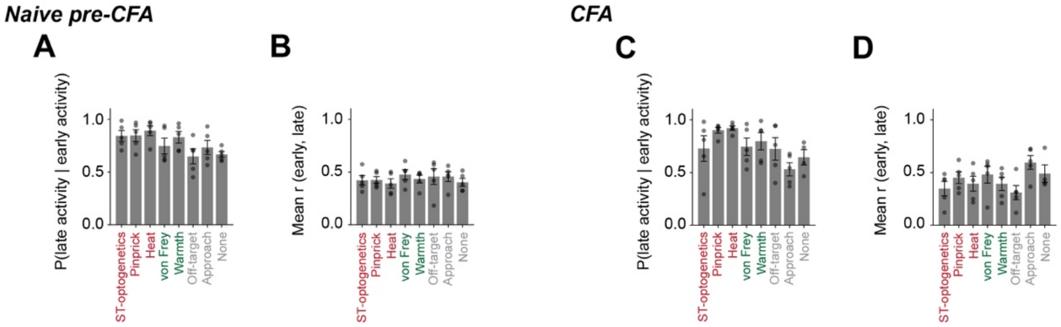


**Supplementary Figure 10 (relates to Fig. 4C). Early–late activity coupling.** (**A, C**) Probability of late activity among neurons classified as early-active, shown for each stimulus. (**B, D**) Mean Pearson correlation between early and late activity, calculated per stimulus and mouse. Bars show mean ± SEM across five mice. These plots show that early and late activity were coupled across stimuli  both before and after CFA.

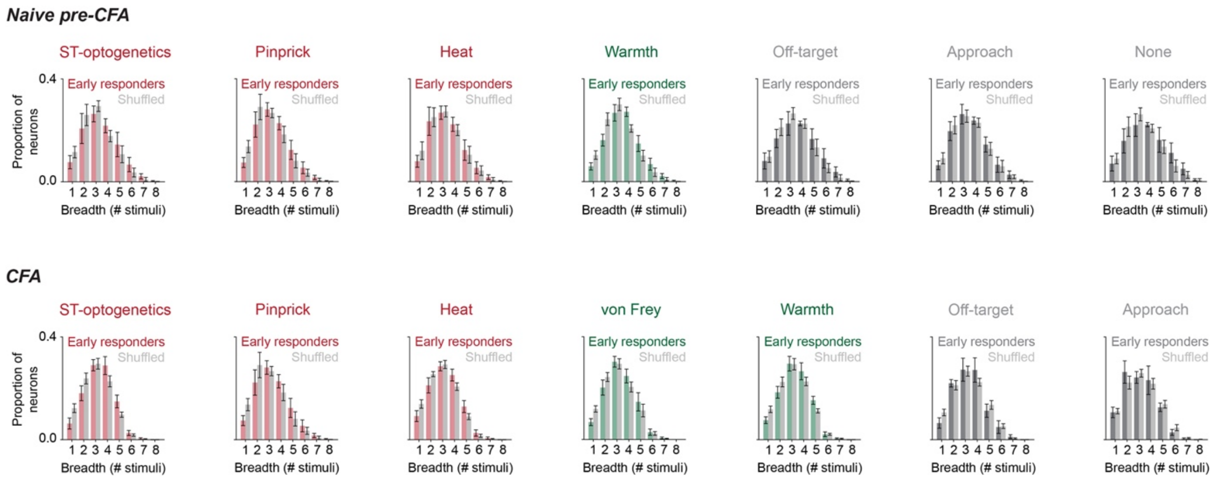


**Supplementary Figure 11 (relates to Fig. 4F). Response breadth across stimuli before and after CFA.** Per-stimulus response breadth of early-responsive neurons. Mean ± SEM across five mice before and after CFA

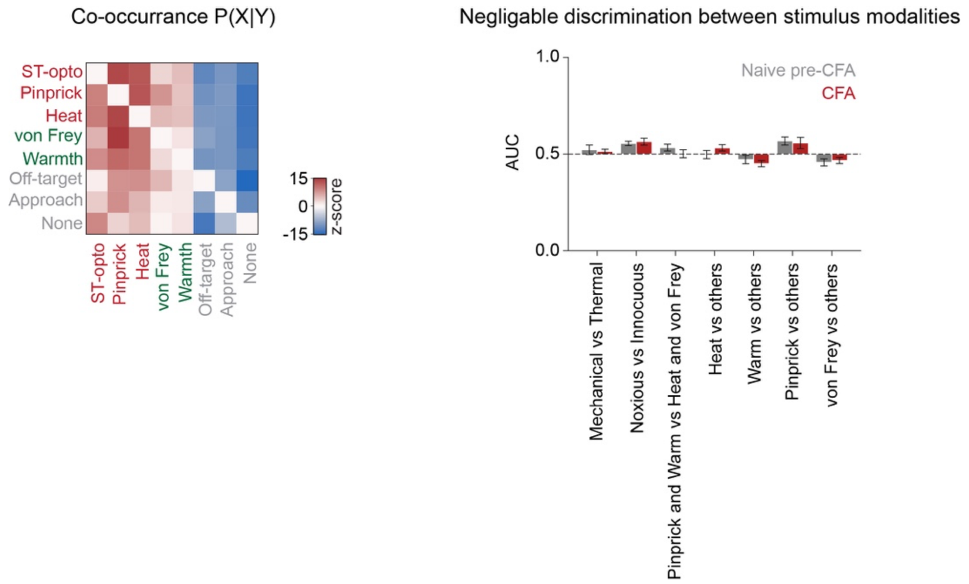


**Supplementary Figure 12 (relates to Fig. 4G).** Co-occurrence and stimulus modality discriminability of early-responsive populations. Left, co-occurrence of early-responsive populations across eight stimulus types. Values show P(B|A), *z*-scored against a response-breadth-preserving label-shuffle null. Right, mouse-mean ROC AUC for stimulus contrasts using early single-neuron activity as the classifier score. AUCs remained close to chance, indicating weak single-neuron stimulus discriminability before and after CFA. All mean ± SEM ($n$ = 5).

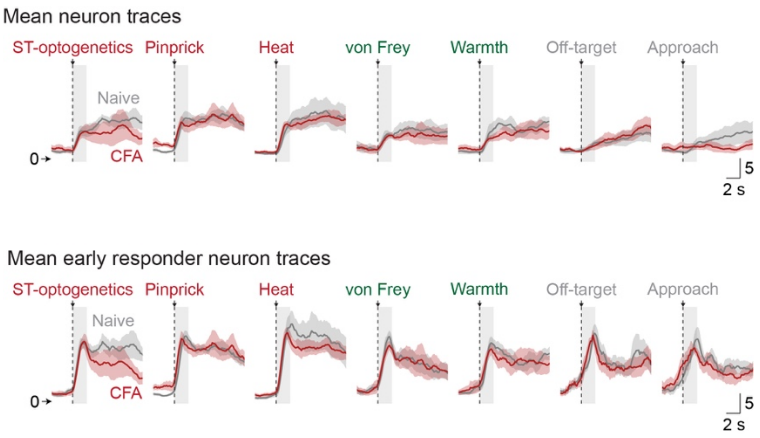


**Supplementary Figure 13 (relates to Fig. 4I).** Mean event-activity traces across stimuli before and after CFA. *Top*, mean neural activity traces for each stimulus, averaged across all trials and neurons within mouse and then across mice. Dashed vertical line marks stimulus onset and grey band marks the early response window. Mean ± SEM ($n$ = 5). *Bottom*, same as top, restricted to neurons classified as early-responsive to each stimulus.

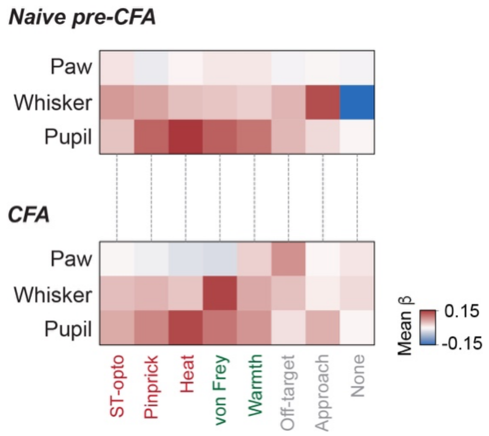


**Supplementary Figure 14 (relates to Fig. 4L). Ridge coefficients across neurons, stimuli, and pain state.** Mean standardized ridge-regression coefficients for models predicting early neural activity from paw distance, whisking, and pupil size. Rows show behavioral predictors and columns show stimuli. Color indicates mean coefficient magnitude and sign.

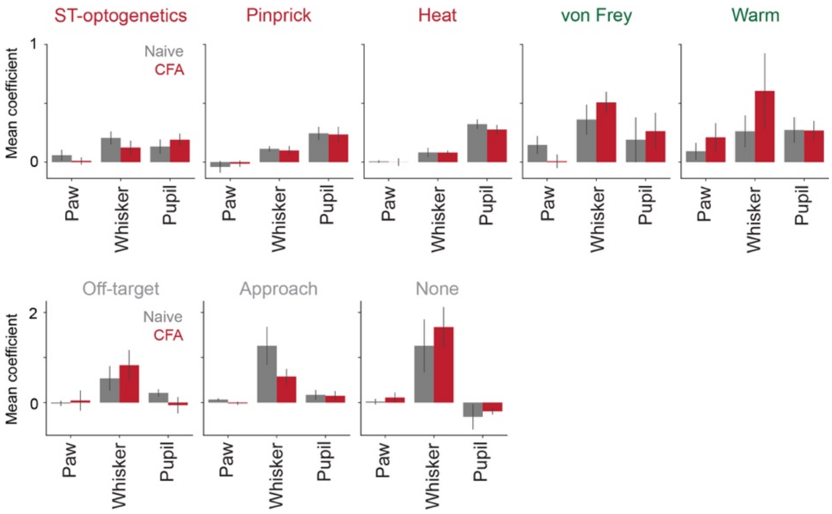


**Supplementary Figure 15 (relates to Fig. 4M). Mean ridge coefficients by stimulus and condition.** Mean standardized ridge-regression coefficients for paw distance, whisking, and pupil size, shown for each stimulus before and after CFA. Bars show mean across five mice and error bars show SEM. Whisking and pupil coefficients were larger than paw coefficients across many stimulus conditions.

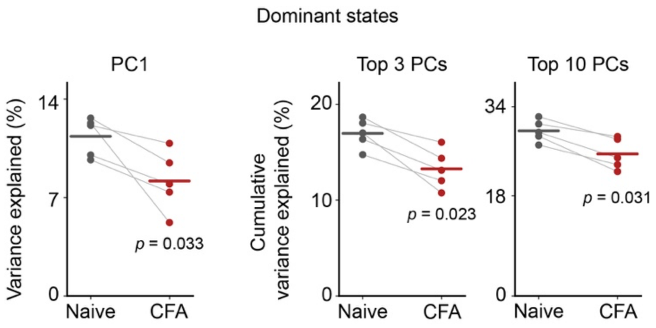


**Supplementary Figure 16 (relates to Fig. 5G). Eigenspectrum.** Variance explained by PC1, cumulative variance explained by the top 3 PCs, and cumulative variance explained by the top 10 PCs before and after CFA. Dots show mice, paired lines connect conditions, and horizontal bars indicate means. *p* values are from exact within-mouse CFA relabeling tests, one-tailed for CFA < Normal, for five mice.

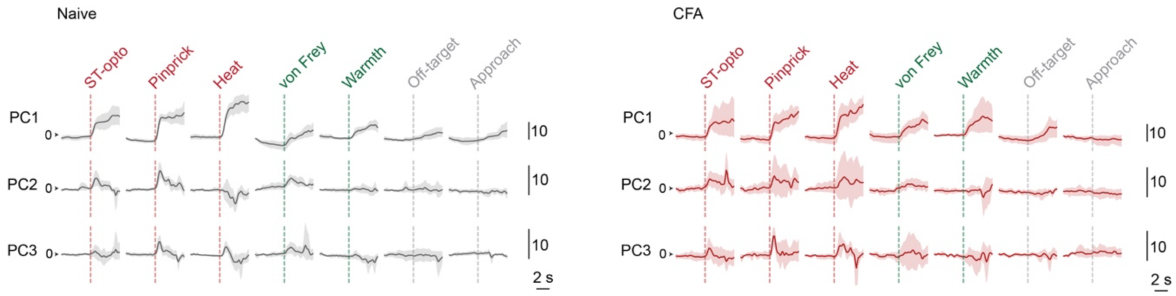


**Supplementary Figure 17 (relates to Fig. 5G). Stimulus-aligned PC traces before and after CFA.** Mean PC1–PC3 time courses for stimuli before and after CFA. Traces show mean ± 95% bootstrap CI across five 5 mice. Dashed lines are stimulus onset.

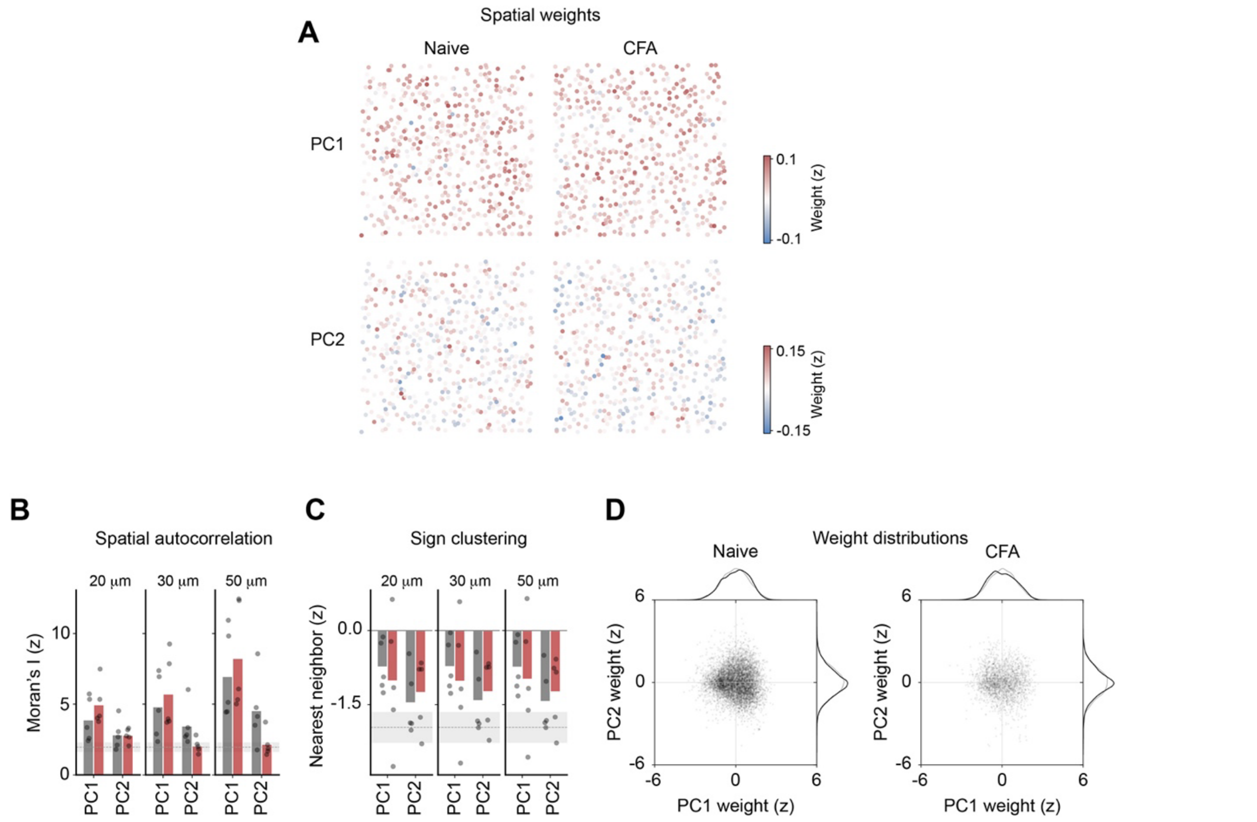


**Supplementary Figure 18 (relates to Fig. 5G). Spatial structure of PC weights.** (**A**) Per-ROI PC1 and PC2 loadings plotted at the Suite2p ROI centroid for one representative session per condition from the same mouse. (**B**) Moran's I of PC loadings (spatial autocorrelation), *z*-scored against 1,000 spatial-permutation nulls, at three inverse-distance scales per condition. Grey shading and dashed line mark the $z = 1.96$ significance threshold ($p = 0.05$ against spatial permutation null). Mean across $n = 5$ mice, with individual mice shown. (**C**) Same-sign nearest-neighbor *z* (distance between nearest-neighbor ROI pairs with loadings of the same sign, *z*-scored against the spatial-permutation null), at the same three scales, per condition. Grey shading is null 95% range. (**D**) Bivariate PC1/PC2 loading distributions across all neurons, separately for conditions. PC1 weights showed significant spatial autocorrelation consistent with smooth spatial gradients, but no evidence of discrete same-sign clustering. This indicates that the state axis was organized as a continuous gradient across the population rather than as distinct functional patches.

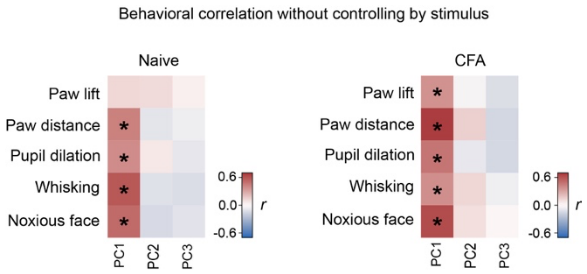


**Supplementary Figure 19 (relates to Fig. 5D,I). PC–behavior correlations without stimulus control.** Pearson correlations between PC scores and behavioral scalars, calculated across all trials pooled across stimuli within each mouse, then averaged across mice. Asterisks mark correlations significantly different from zero by exact sign-flip test across mice, $p < 0.05$. n = 5 mice.

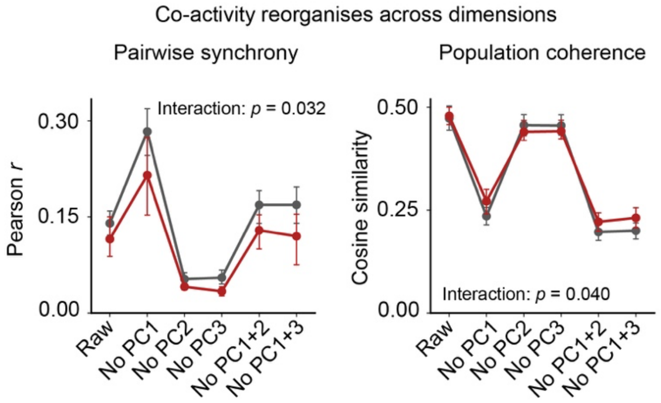


**Supplementary Figure 20 (relates to Fig. 5K,L). Co-activity after PC removal.** Pairwise Pearson synchrony and population cosine coherence after removing PC1, PC2, PC3, PC1+2, or PC1+3 from population activity. Mean ± SEM for five mice. *p* values test the condition and PC-removal interaction using exact within-mouse CFA relabeling.

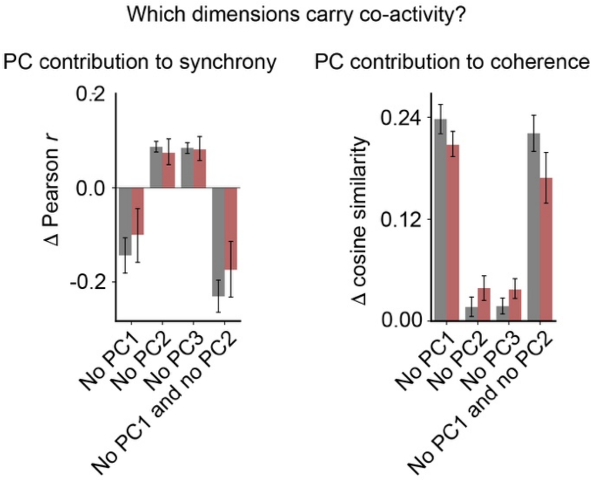


**Supplementary Figure 21 (relates to Fig. 5L). PC contributions to co-activity.** Change in pairwise Pearson synchrony or cosine coherence after removing each PC or PC pair, defined as raw minus PC-removed co-activity. Positive values indicate that the removed component contributed positively to co-activity. Mean ± SEM for five mice.

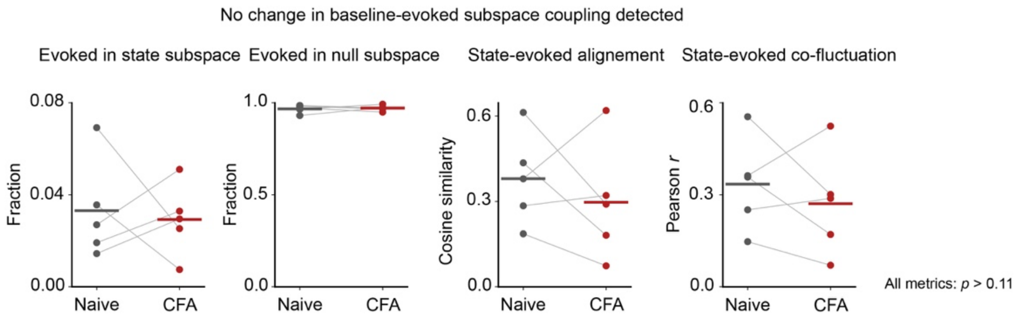


**Supplementary Figure 22 (relates to Fig. 5L). Baseline–evoked subspace coupling.** State-window (−10 to −5 s) and evoked-window (−2 to +3 s) subspace coupling before and after CFA. Metrics quantify the fraction of evoked variance in the state subspace, the complementary null-space fraction, PC1 state-evoked alignment, and state-evoked co-fluctuation. Exact within-mouse CFA relabeling detected no significant changes. n = 5 mice.

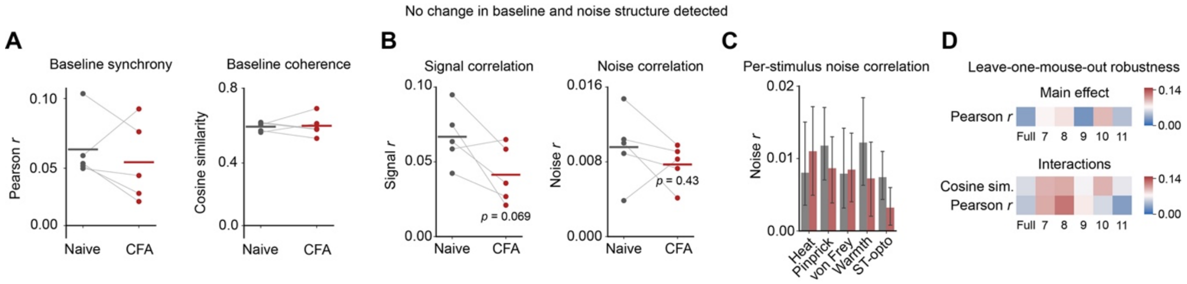


**Supplementary Figure 23 (relates to Fig. 5K,L,M). Baseline and noise structure controls.** (**A**) Baseline synchrony and coherence, evoked signal and noise correlations, per-stimulus noise correlations, and leave-one-mouse-out robustness for the synchrony and coherence analyses. Baseline synchrony and baseline coherence over the state window (−10 to −5 s) did not change significantly across *n* = 5 mice (exact within-mouse CFA relabeling). (**B**) Pairwise signal correlation and noise correlation over the evoked window, computed on the five stimuli (heat, pinprick, von Frey, warmth, ST-opto). (**C**) Per-stimulus noise correlation breakdown across five stimuli. Mean ± SEM (*n* = 5). (**D**) Leave-one-mouse-out robustness show condition main-effect and PC-removal interaction p-values (exact within-mouse CFA relabeling, $4^5$ = 1,024 or $4^4$ = 256 permutations when each mouse is dropped in turn) for the pairwise Pearson synchrony and cosine coherence analyses of Fig. 5L and Supplementary Fig. 21,22. This indicated that no single mouse drove the effect. Together, this showed that the co-activity reorganization reported in Fig. 5 reflected a redistribution across dimensions rather than a change in baseline co-activity, noise structure, or any single-mouse effect.

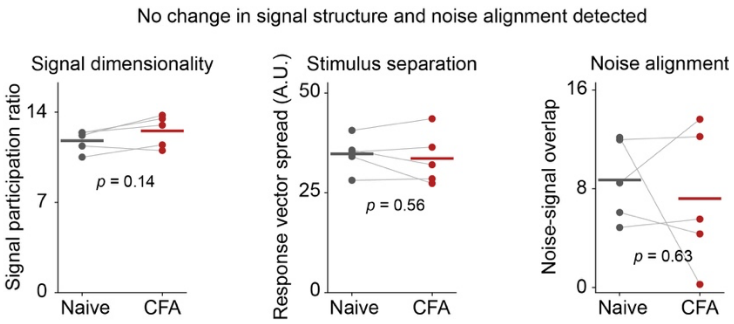


**Supplementary Figure 24 (relates to Fig. 5N). Signal geometry controls.** Signal participation ratio, stimulus mean position spread, and noise-signal overlap from the five-stimulus (heat, pinprick, von Frey, warmth, ST-opto; trial counts balanced per session) analysis. Stimulus mean positions were computed from mean activity over 0 to 1.5 s post-stimulus, with trial counts balanced per session. Dots show mice, paired lines connect conditions, and horizontal bars indicate means. *p* values are from exact within-mouse CFA relabeling tests. *n* = 5 mice. None of the three changed significantly, indicating that the reduction in stimulus contrast alignment (Fig. 5N) was not explained by a change in the magnitude or the noise structure of the signal subspace.

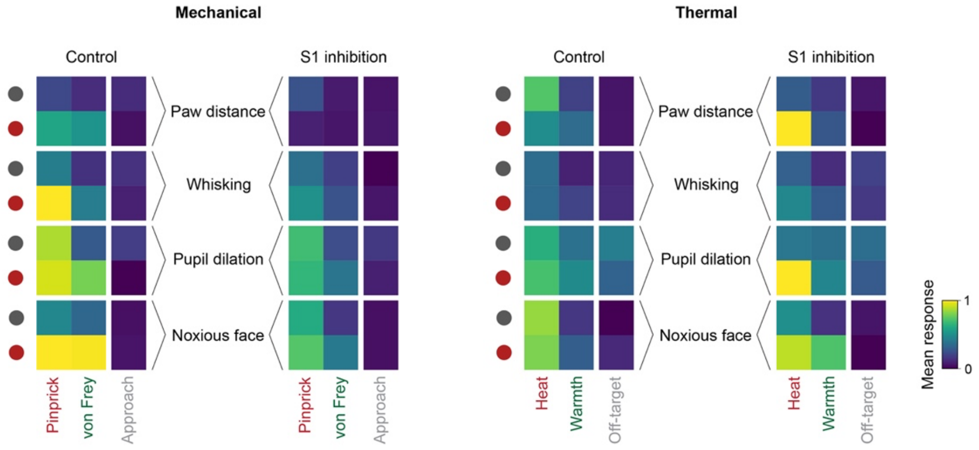


**Supplementary Figure 25 (relates to Fig. 6). Per-condition and per-treatment response heatmaps for the inhibition cohort.** Mean behavioral responses across the four condition × treatment blocks of the S1 inhibition experiment. Grey circles indicate before CFA and red circles indicate after CFA. Cell values are per-mouse mean responses, min–max scaled within each behavioral feature across both treatment conditions. *n* = 6 mice. Mechanical control trials showed CFA-enhanced responses that were reduced during S1 inhibition. Heat responses were already high before CFA and increased little after it, limiting the dynamic range over which sensitization could be detected. S1 inhibition did not reduce heat- or warmth-evoked responses. We note that heat responses appeared, if anything, larger during inhibition. Warmth responses were lower at baseline, allowing CFA-enhanced responses to be detected, and these were not reduced by S1 inhibition.

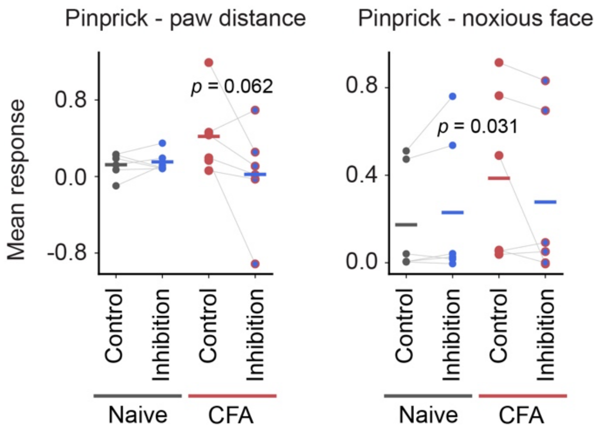


**Supplementary Figure 26 (relates to Fig. 6). Pinprick responses during S1 inhibition.** Per-mouse pinprick responses. *Left*, paw distance. *Right*, noxious-face score. p-values are exact one-tailed sign-flip permutation tests on the interaction. *n* = 6 mice.
